# Perturbational single-cell profiling of patient tumors defines lineage- and context-specific programs of innate immune resistance

**DOI:** 10.64898/2025.12.09.693245

**Authors:** C. Perry, A. Frey, Y. Fei, J. Wang, P. Costa, H. N. Amin, S. Ghadermarzi, D. Levine, W. Tong, A. Koda, M. Mackie, M. He, Y. Nie, K. Clulo, F. Ouerghi, J. Wei, T. Cordero Dumit, M. Yaskolko, M. Ding, A. Caldera, O. Kyrysyuk, L. Lum, J.W. Allen, W. Guo, A. Naqash, A. Elliot, A. Vanderwalde, M. Capelletti, T. Adeyelu, S.T. Barry, M. Hugaboom, A. Bacchiocchi, H. Kluger, M. Bosenberg, A. Iwasaki, D. Braun, J. Clune, D. van Dijk, K. Olino, J.J. Ishizuka

## Abstract

Despite promise in preclinical models, most immuno-oncology drug candidates fail in clinical trials. These failures reflect limitations in our ability to directly model the response of human tumor and immune cells to immunotherapies. To address this gap and test the effect of innate immune agonists, we developed PERCEPT, an approach that uses *ex vivo* perturbational single-cell RNA sequencing to compare the response of immunomodulatory treatments with unstimulated controls directly in patient samples. Using PERCEPT, we tested cytokines and innate immune agonists in melanoma and Merkel cell carcinoma (MCC) and identified the dsRNA mimetic, RIG-I agonist, Stem Loop RNA (SLR) 14 as a powerful inducer of anti-viral states and enhancer of T cell activation. We compared transcriptional ‘responder’ and ‘non-responder’ patient samples and identified midkine (MDK), a multifunctional cytokine, as a potent repressor of IFN signaling in both tumor and immune cells. MDK expression dampened MHC-I presentation in human tumor cells and reduced activation of antigen-presenting cells, disrupting tumor immunity at multiple levels. In contrast to prior studies, we identified MDK as specifically enriched in neuroendocrine cancers such as MCC and small cell lung cancer compared with melanoma, suggesting the importance of lineage- and context-specific targeting. Our results demonstrate the utility of high-dimensional controlled perturbation of patient samples to identify mechanisms of innate immune response and resistance and demonstrate an actionable path towards clinical development of MDK-inhibiting therapies including FDA-approved ALK inhibitors in neuroendocrine cancers.

## Main

The success of immunotherapy in cancer has led to the preclinical development of thousands of novel immunotherapies with over one thousand approaches tested in clinical trials^1^. However, the vast majority of these therapies have failed in clinical development^2^. A principal reason for failure is the lack of translation of efficacy signals from preclinical murine and cell culture model systems to clinical trials, and the underlying limitations of these systems in modeling complex and heterogeneous patient tumor-immune microenvironments, which are shaped by months or years of tumor-immune co-adaptation^3^. The experience of multiple negative Phase III clinical trials with DMXAA, ultimately shown to not stimulate STING in humans as it does in mouse models, underscores the perils of pre-clinical translation^4^. High-dimensional studies of human tumors, including single-cell RNA sequencing and spatial approaches, have helped to reveal the complexity of patient tumor-immune interactions. However, these approaches have most frequently been used to model tumor and immune states prior to, or significantly after perturbation, and have only sparingly been used with high-dimensional approaches to directly assess the response of therapies using primary tumor and immune cells^5–10^. This gap is notable, given that immuno-modulators frequently rewire tumor-immune interactions in emergent and unexpected ways. Similarly, although the development of generative artificial intelligence and large language models offer the potential to improve predictions of clinical efficacy, these models currently lack training data that maps the transcriptional state response of complex, interacting immune cell mixtures to perturbations.

Interferon (IFN)-stimulating therapies that induce viral mimicry, including STING agonists, oncolytic viruses, and double-stranded RNA (dsRNA) mimetics, represent a prominent class of novel drugs in which promising preclinical studies have underperformed in clinical trials^11,12^. This lack of development is striking given the strong overlap of these pathways with the recruitment and activation of anti-tumor immune populations, including classical dendritic cells, CD8+ T cells, NK and gamma-delta T cells, as well as extensive supportive evidence from murine studies^13–15^. For example, the RIG-I agonist Stem Loop RNA (SLR)14, a 14-base-pair double-stranded RNA hairpin, can sensitize “cold” tumors to ICB via direct intratumoral injection, leading to responses at distal tumor sites and immunologic memory^1^. To better define the mechanisms that regulate the response of human tumor microenvironments to immunotherapies and nucleic acid sensor agonists, we developed PERturbational sequencing of immune Co-culture from *Ex vivo* Patient Tumors (PERCEPT), an approach that performs single-cell RNA sequencing of patient tumor and immune cells following multiplexed therapeutic perturbation. Critically, therapeutic perturbations are compared with unstimulated controls, enabling the application of advanced causal inference approaches and the isolation of true response from confounder variation among heterogenous patient samples^2^. PERCEPT builds on prior approaches demonstrating prediction of clinical response to immunotherapy with *ex vivo* tumor perturbation, but addresses intratumoral heterogeneity using homogeneous mixtures of tumor and immune cells and response complexity by integrating scRNAseq with flow cytometry and multiplexed cytokine analysis^9^.

We hypothesized that PERCEPT would enable the identification of cell type and tumor type-specific mechanisms of treatment response and resistance in human tumors. To test this hypothesis, we applied PERCEPT to an array of checkpoint inhibitors, cytokines, and innate immune agonists, including SLR14 in melanoma and Merkel cell carcinoma (MCC), two lethal skin cancers in which these therapies are under investigation^16^. We detected a range of cell and tumor-type-specific responses, including a marked activating response within the T cell compartment to SLR14, in contrast to the minimal response we observed to STING agonism. In several patient samples, the response to nucleic acid agonism was muted or nearly absent.

Applying a causal inference framework^17^, we identified the cytokine midkine (MDK) as an innate immune checkpoint limiting IFN response and MHC-I upregulation and specifically upregulated in MCC compared with melanoma, where it has most intensively been studied previously. MDK expression was inversely correlated with IFN response to stimulation with a range of perturbations. We validated the potential for MDK to suppress IFN responses and MHC-I expression using CRISPR in both human tumor cell lines and primary immune cells, including monocytes and monocyte-derived dendritic cells. Moreover, we found that this mechanism of immunosuppression applied equally to small cell lung cancer (SCLC), another neuroendocrine tumor with high MDK expression. Our results are consistent with recent work, suggesting that MDK can suppress dendritic cell activation by modulating NF-kB and STAT3 signaling in mouse models of MDK-expressing melanoma, but integrate patient tumor-immune response information to identify a distinct translational development path in human neuroendocrine tumors. To this end, we identify MDK expression as a predictor of resistance to immune checkpoint inhibitor (ICI) treatment in large clinical data of SCLC. Together, these findings support the use of high-dimensional perturbational analysis to help identify mechanisms of treatment response and resistance in humans and to identify patient populations likely to respond to novel therapies.

## Results

### Development of perturbational sequencing from patient tumors

To enable perturbational sequencing from patient tumors, we isolated homogeneous single-cell suspensions of tumor-immune mixtures in their native states and proportions directly from fresh melanoma and MCC surgical resections (**Fig. 1a; Fig. S1a**). We treated independent mixture cell cultures for up to 48 hours with a range of immunomodulatory treatments, including standard of care, developmental and novel immunotherapeutics (**Fig. 1b**). Following stimulation and co-culture, we used Fluorescently Activated Cell Sorting (FACS) to isolate live cells and to quantitate immune infiltration, enriching for immune cells up to 50% of the total mixture in tumor samples with low endogenous immune infiltration. To enable multiplexing, we used cell-hashing to barcode treatment conditions and performed scRNAseq along with cytokine bead array analysis on 8-10 simultaneous treatment conditions. As independent controls, we repeated this workflow using healthy donor peripheral blood mononuclear cells (PBMCs). Given that PD-1 axis-targeting ICIs are the standard of care for advanced disease in both melanoma and MCC, we tested PD-1 in all tumor samples as well as PD-1-based combinations with IFNβ, which we selected as a common biological pathway downstream of nucleic acid sensor stimulation. As a control and to determine overall IFN-axis response in samples, we stimulated tumor-immune mixtures with IFNβ and IFNγ. For novel therapies, we focused on a range of nucleic acid sensor agonists, including transfected and non-transfected polyI:C, a synthetic dsRNA agonist that stimulates a range of dsRNA receptors, ADU-S100, a STING agonist that previously failed in clinical development, and SLR14, a novel RIG-I agonist with promising preclinical studies. Because polyI:C is known to trigger both TLR3, which is accessible on cell surfaces and endosomes, as well as MDA5 and RIG-I, which are cytosolic sensors, we compared the effects of transfected and non-transfected polyI:C^1718^ . In total, we recovered 1.8 x 10^5^ cells from nine tumor and three healthy control samples, representing 100 distinct sample x perturbation combinations and thousands of cell type/state x perturbation combinations, depending upon the specificity of cell type and state definition (**Fig. 1c-e, S1b**). To more specifically assess transcriptional response to stimulation across a broad range of cell types, we used the MELD algorithm (MELD)^19^. We first examined high-dose IFNβ as a positive control and observed signatures of response in nearly all samples and cells, although the strength of response varied widely by individual sample and by sample type (**Fig. 1f**). We also observed the induction of IFN signatures in response to a αPD-1 in specific tumor samples MCC2 and MEL4 that had not previously progressed on αPD-1, but no responses to αPD-1 alone in patients’ tumors which had progressed in response to αPD-1 (**Fig. 1g**). We observed corresponding increases in cytokine production by cytokine bead array to observed changes in scRNAseq (**Fig. S1c,d**).

**Fig 1.**
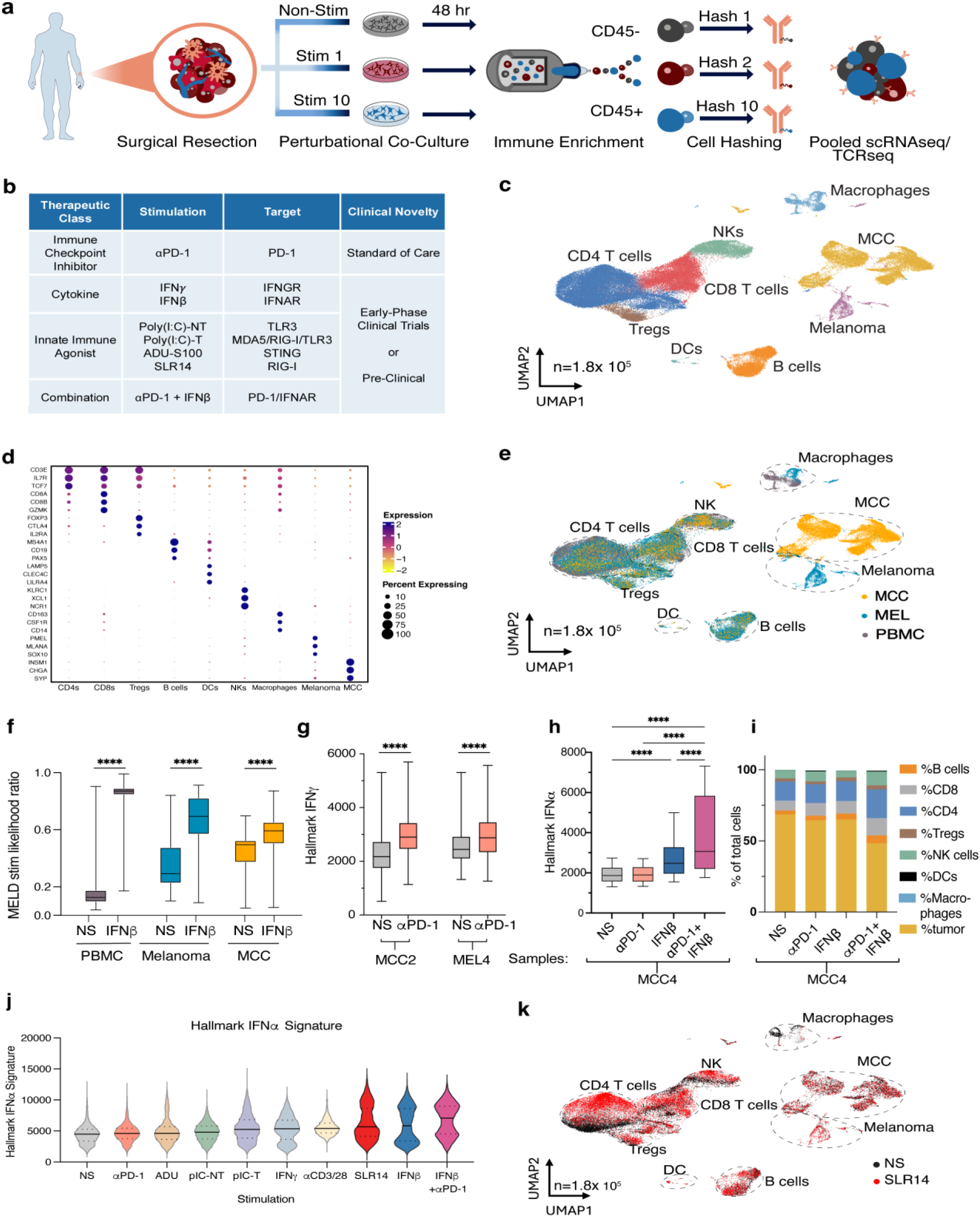
*Ex vivo* single-cell RNA sequencing reveals cell- and tumor type-specific effects of combinatorial innate immune and checkpoint blockade perturbation. (**a**) Study design for *ex vivo* PERCEPT experiments. Tumors were processed *ex vivo* to single-cell suspension, then plated in up to 9 distinct stimulation conditions (**b**) + unstimulated for 42-48 hours, before FACS enrichment of live CD45+ immune cells. Individual treated cells were hashtagged and submitted for 10x single-cell sequencing. (**c**) UMAP of scRNAseq clusters by cellular identity (**d**) as defined by gene expression signature and (**e**) by sample type. (**f**) MELD stimulation likelihood scores to IFNβ by tissue type. (**g**) ssGSEA for hallmark IFNγ response signature of tumor cells increased by αPD-1 for samples MCC2 and MEL4. (**h**) ssGSEA for hallmark IFNγ response signature of tumor cells increased by αPD-1, IFNβ, or both, for sample MCC4. (**i**) % cell type by scRNAseq of sample MCC4 with % tumor cells decreased by the combination of both αPD-1 and IFNβ. (**j**) ssGSEA of hallmark canonical IFNα signature with the most marked response to SLR14, IFNβ, and IFNβ+PD-1. (**k**) UMAP of all cells after SLR14 (red) compared to non-stimulated (black).

Moreover, we were able to detect cooperative increases in IFN signaling and decreased percentage and numbers of tumor cells when IFNβ and αPD-1 were combined for the sample MCC4, which demonstrated no response to αPD-1, and had progressed in response to clinical αPD-1 treatment (**Fig. 1h,i, S2a,b**), suggesting the potential for PERCEPT to be applied to dissect combinatorial contributions to signaling. We observed synergistic responses to IFNβ and αPD-1 in melanoma tumor cells, as well as NK and T cells from MCC tumor samples, largely reflecting cell-type-specific synergy from MCC2 and MCC4 (**Fig. S2c,d**)^17^. Comparing the effect of type I IFN-inducing therapies across all tumor-immune and healthy donor samples, we noted a progression of effects, with αPD-1 and ADU-S100 inducing mild IFN-induction except in exceptional samples, but SLR14, IFNβ, and the combination of IFNβ and αPD-1 inducing the most potent effects when averaged over all cell types (**Fig. 1j**).

### T cell programming by SLR14

Of the novel agents, SLR14 treatment conferred particularly strong treatment effects across all cell types (**Fig. 1k**). Further, although SLR14 strongly induced canonical interferon-stimulated genes (ISGs) it also triggered expression of a distinct subset of genes including genes associated with T cell activation and cytotoxicity including *CCL5*, *PRF1* and *CCR7* (**Fig. S2e,f**). Indeed, SLR14 induced distinct activation states in CD8+ T cells (**Fig. 2a, S2g**), CD4+ T cells (**Fig. 2d, S2h**), and NK (**Fig. S2i**) cells compared to even supraphysiologic levels of IFNβ, with a marked upregulation of both STAT and NF-kB regulons. Following SLR14 stimulation, PERCEPT revealed shifts in the transcriptional phenotypes of stem-like progenitor and terminally differentiated T cell populations^20–22^, including the induction of antiviral CD4+ and CD8+ populations not observed in control conditions^23^ (**Fig. 2c**). In tumor cells, NK cells and T cells, SLR14 stimulation induced expression of canonical IFN-stimulated genes (**Fig. 2d, bottom**). In CD8+ T cells, however, we observed specific programs of activation and survival including *CD69*, *IL15RA and IL2RG* and decreased expression of the exhaustion markers *TOX* and *PDCD1* (**Fig. 2d, top**)^22^. CD4+ T cells were also reshaped by SLR14 exposure, with decreased expression of *TOX*, *FOXP3*, *TGFB1*, and *LGALS1*, consistent with decreased regulatory T cell polarization. Both CD8+ T cells and CD4+ T cells expressed increased levels of B-cell activating factor (BAFF, *TNFSF13B*), suggesting the potential for supportive interactions between B and T cells^23,24^. Given the transcriptional evidence for T cell activation and reprogramming following SLR14 treatment, we hypothesized that SLR14 may shift the T cell activation threshold. To test this hypothesis, we treated healthy donor PBMCs with SLR14 or control for 48 hours prior to activation with αCD3/CD28 beads (**Fig. 2e, S2j**). We observed increased production of IFNγ in both the CD4+ and the CD8+ compartments in SLR14-treated samples compared to lipofectamine-treated controls.

**Fig 2.**
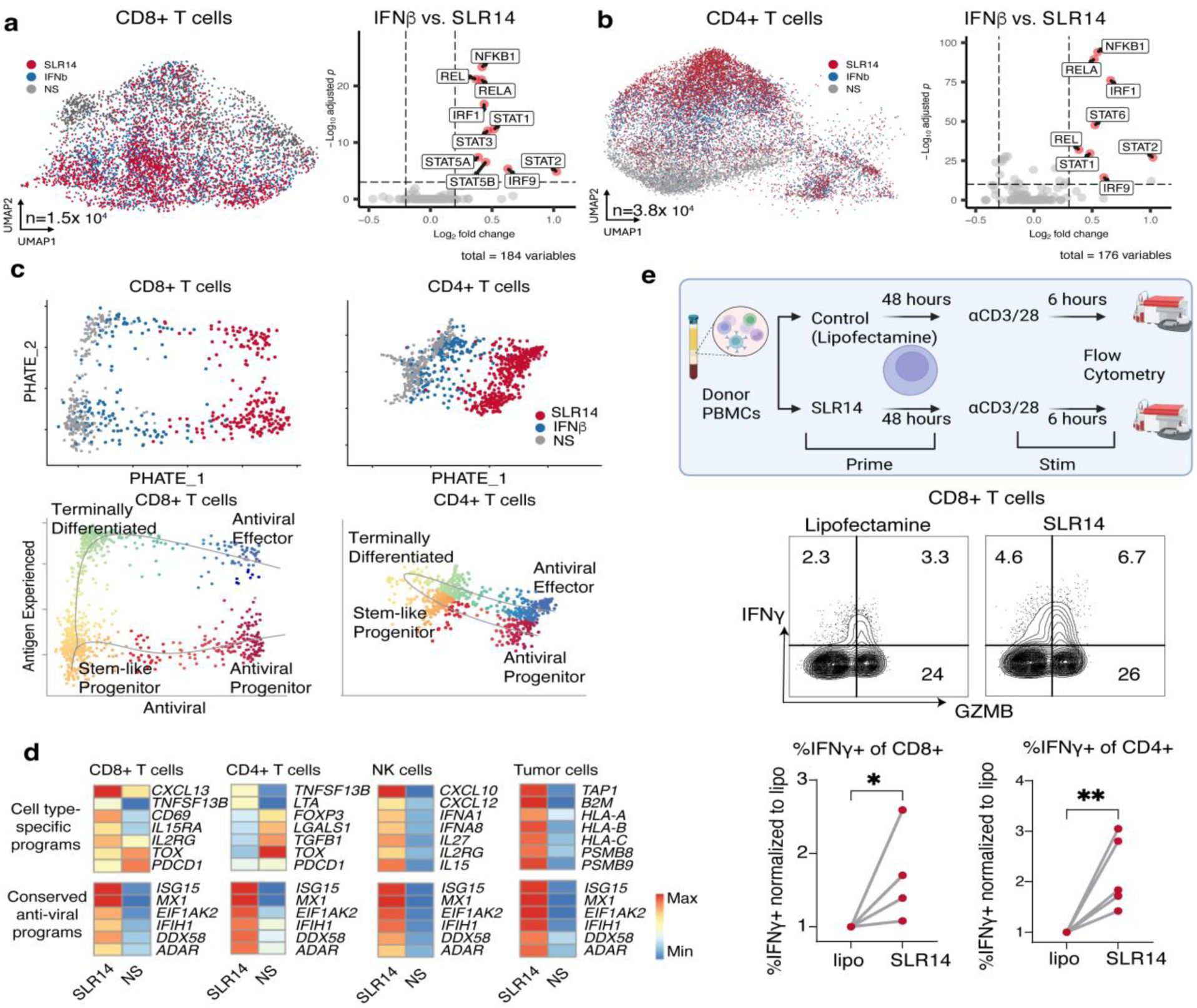
RIG-I priming enhances T cell response in tumor-immune co-culture via NF-kB signaling. (**a**) UMAP and volcano plot of VIPER regulons of CD8 and (**b**) CD4 T cells after SLR14 and IFNβ. (**c**) PHATE plot of CD8 and CD4 T cells stimulated with SLR14 and IFNβ from sample MEL5. (**d**) Specific gene programs from CD8, CD4, NK and tumor cells from sample MEL5. (**e**) PBMCs were isolated and primed with SLR14 or lipofectamine control for 48 hours as per *ex vivo* tumor protocol, before 6-hour stimulation and αCD3/28 stimulation in Brefeldin A to measure intracellular IFNγ production. P < *0.05, **0.01, from t-test between lipofectamine control and SLR14.

### Defining response and resistance patterns in patients

Having identified IFNβ and SLR14 as the two most potent inducers of transcriptional tumor-immune response in our cohort, we next sought to identify the patterns and determinants of response to these two agents. To quantify response across diverse cell types and samples, we again used MELD^19^. Although responses to IFNβ and SLR14 demonstrated sample-specific variation, with some samples responding more strongly to one treatment than to another, responses to the two therapies correlated overall within our cohort (**Fig. S3a**). Further, we identified a subset of samples in which neither tumor nor immune cells demonstrated a robust response to either therapy. Specifically, we identified six tumor samples in which robust transcriptional responses were noted to either IFNβ or SLR14 Responders (R), in contrast to three Non-Responders (NR) samples with minimal response to either treatment (**Fig. 3a, 3b**). Comparing all samples, we noted a significantly greater response to both IFNβ and SLR14 in melanoma compared to MCC (**Fig. 3c**). Following stimulation with IFNβ or SLR14, Responder samples upregulated signatures of IFN signaling, allograft rejection, inflammatory response and NF-kB signaling compared to Non-responders (**Fig. 3d**). Given the enrichment of NR samples in MCC compared to melanoma, we sought to identify the patterns of signaling and potential mechanisms underlying MCC non-response. To this end, we employed causal independent effect module attribution + optimal transport (CINEMA-OT), a causal inference framework that uses optimal transport matching to isolate confounder from response variation in matched samples^17^. Using CINEMA-OT, we generated a Treatment Effect Space (TES) in which to compare MCC R and NR samples (**Fig. 3e**) and derived the genes that were most notably differential within this space (**Fig. 3f**). This analysis highlighted the multifunctional cytokine midkine (*MDK*) as the most significantly upregulated gene in the TES of MCC NR compared to MCC R samples. Consistent with CINEMA-OT analysis, MDK expression was higher overall in MCC NR than MCC R tumor cells (**Fig. 3g**)^17^. Notably, MDK expression was also significantly higher in MCC than melanoma cells. Because MDK is pleiotropic, with numerous potential ligand-receptor pairs, we used CellChat to identify putative drivers of signaling in our cohort^11,16,25,26^.**Fig. 3h, S3b** CellChat identified MCC, but not melanoma tumor cells as a primary source for MDK signaling (), with tumor cell autocrine signaling via nucleolin or PTPRz1 and putative signaling to myeloid cells including monocytes and DCs via NOTCH2, nucleolin and/or LRP1 receptors (**Fig. 3h, S3c**).

**Fig 3.**
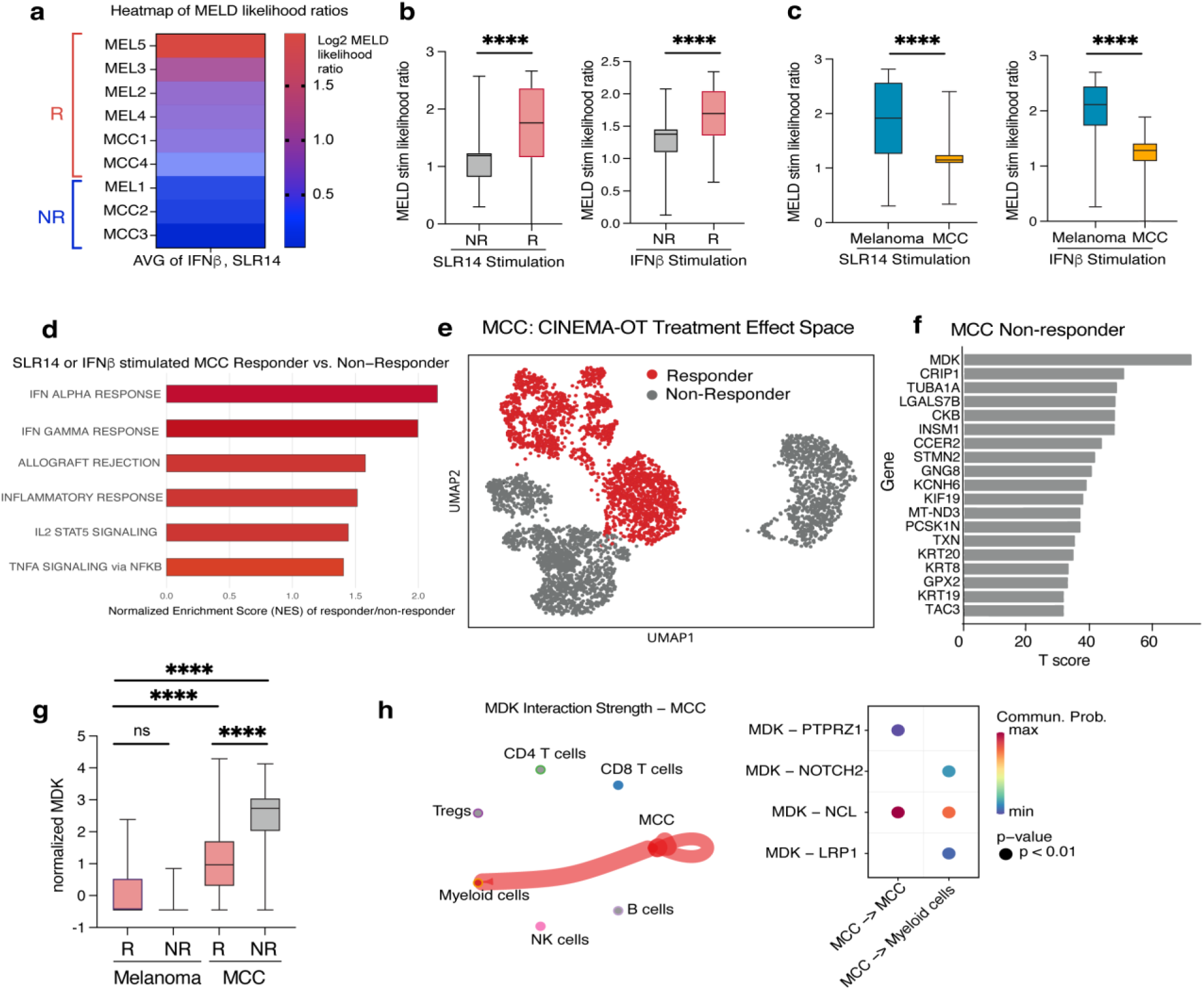
A multiplexed single-cell perturbational platform identifies midkine associated with suppressed IFN responses in patient samples. (**a**) PERCEPT samples’ likelihood of response to SLR14 or IFNβ by MELD score was measured, and samples were categorized into Responder (R) by average log2 MELD score of SLR14 or IFNβ >0.5 and Non-Responder (NR) MELD <0.5. MELD score of each single cell from (**b**) R vs. NR samples and (**c**) melanoma vs. MCC samples. ****=p < 0.001 by Mann-Whitney test. (**d**) R and NR MCC samples stimulated with SLR14 or IFNβ were compared by ssGSEA of Hallmark gene sets, and significant (p>0.05) pathways were plotted by Normalized Enrichment Score (NES). (**e**) CINEMA-OT was performed, and the treatment effect space of MCC samples was compared R vs. NR, (**f**) with significantly increased genes (t-score>30) in the treatment effect space. (**g**) MDK scRNAseq read count of each tumor cell from melanoma and MCC samples by R vs. NR. ****=p < 0.001 by two-way ANOVA with Sidak correction for multiple comparisons. (**h**) CellChat analysis of the MDK signaling pathway from MCC samples by cell type and candidate MDK receptors (highest 4%).

### Validation of MDK effects in neuroendocrine tumor cells

Given that the primary prior work in the field identified midkine as a regulator of response to immune checkpoint inhibition in melanoma, we were surprised to observe a stronger effect in MCC than in melanoma patient samples. We hypothesized that MDK signaling may be enriched in MCC due to its neuroendocrine differentiation and that its inhibition may be of particular translational importance in neuroendocrine tumors such as MCC and Small Cell Lung Cancer SCLC)^27^. To test this hypothesis, we compared the expression of MDK in bulk RNAseq datasets from melanoma, MCC and SCLC. Although MDK was expressed in certain samples in all three diseases, we identified that MCC and SCLC had significantly greater expression of MDK than melanoma (**Fig. 4a**), which was confirmed in comprehensive expression profiling across a wide variety of patient tumor types (**Fig. S4a**). Consistent with our PERCEPT profiling, scRNAseq from patient tumors with SCLC confirmed that tumor cells are the predominant source of midkine (**Fig. 4b**).

**Fig 4.**
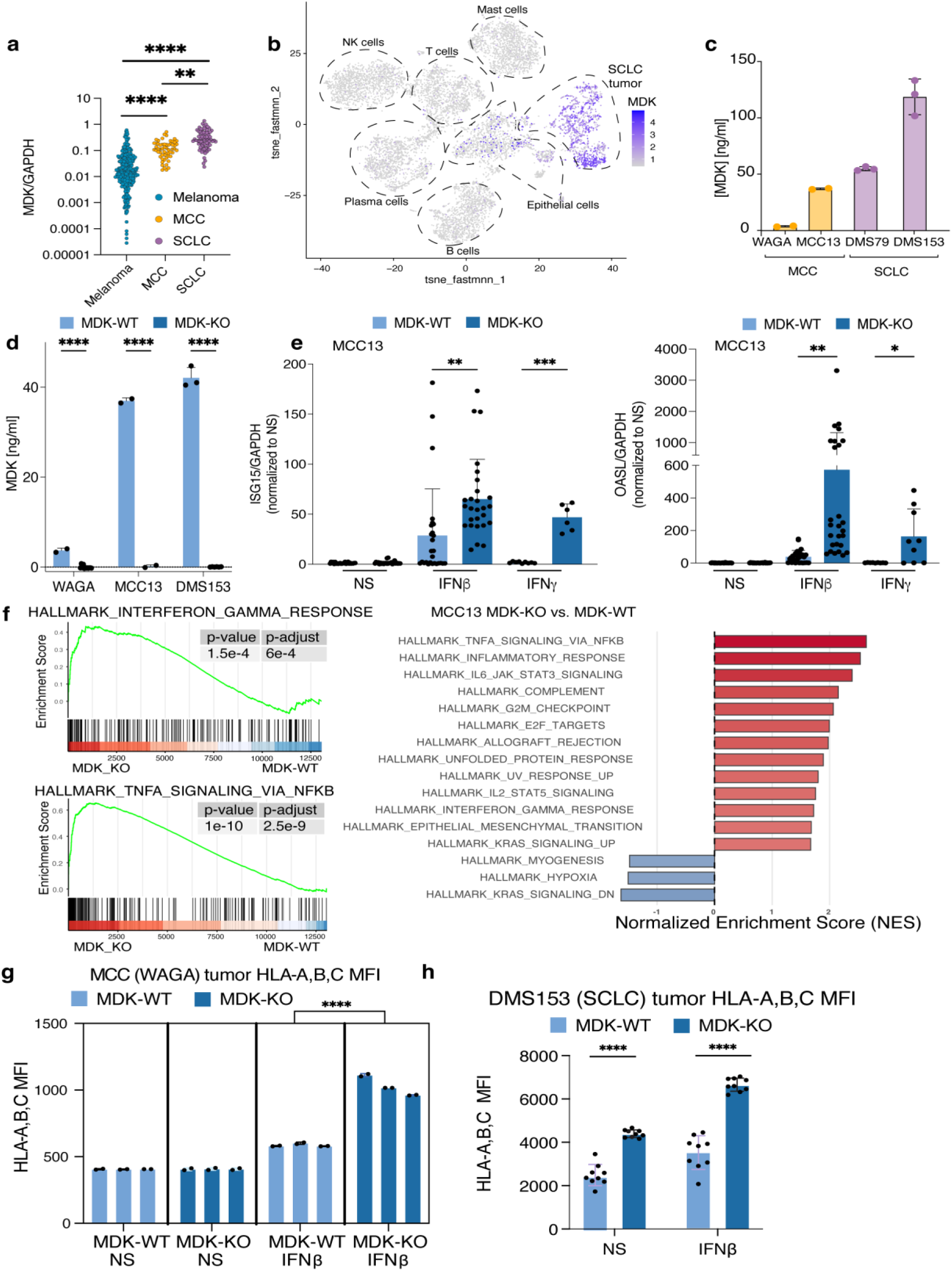
Midkine (MDK) suppresses IFN response and HLA class I upregulation in MCC and SCLC. (**a**) Bulk RNAseq MDK/GAPDH of patient samples: 473 melanoma^29^, 54 MCC^30^, 86 SCLC^31^. **=p < 0.01, ****=p < 0.001 by two-way ANOVA with Sidak correction for multiple comparisons. (**b**) t-SNE-fastMNN of scRNAseq of SCLC patients with MDK expression by read count. (**c**) MDK ELISA of MCC and SCLC cell line supernatant. (**d**) MDK ELISA of indicated MDK-WT and MDK-KO cell line supernatant. (**e**) MCC13 MDK-WT and MDK-KO treated with 1000 IU/ml IFNβ or 1000 U/ml IFNγ for 24 hours prior to RNA isolation and qPCR for ISG15/GAPDH and OASL/GAPDH, normalized to non-stimulated (NS) **=p < 0.01, ***=p < 0.005 by two-way ANOVA with Sidak correction for multiple comparisons. (**f**) ssGSEA of p-adjust<0.05 by Normalized Enrichment Score (NES) of RNAseq of MCC13 MDK-WT and MDK-KO treated with 1000 IU/ml IFNβ for 24 hours. (**g**) WAGA (MCC) or (**h**) DMS153 (SCLC) MDK-WT and MDK-KO tumor cells treated with 1000 IU/ml IFNβ vs. NS for 24 hours prior to flow cytometry of HLA A,B,C on live cells. ****=p < 0.001 by two-way ANOVA with Sidak correction for multiple comparisons. Data representative of three independent experiments.

To enable experimental validation of our patient sample observations, we sought to confirm the high levels of MDK expression in MCC and SCLC cell lines. We observed prominent expression of MDK in MCC cell lines (WAGA, MCC13) and even higher expression in SCLC cell lines (DMS79, DMS153) (**Fig. 4c**). To interrogate the effect of MDK loss on IFN response and tumor immunity, we used CRISPR-Cas9 to genetically delete *MDK* in the MCC cell lines WAGA and MCC13 (**Fig. 4d**). Consistent with PERCEPT results from patient samples, deletion of *MDK* significant enhanced the expression of ISGs including ISG15 and OASL following stimulation with either IFNβ or IFNγ (**Fig. 4e**). In addition to altered ISG response and consistent with prior evidence of MDK as a tumor growth factor, we observed slower growth of WAGA and MCC13 following deletion of *MDK* (**Fig S4b**^26^. To assess the transcriptional rewiring associated with MDK deletion, we performed RNAseq of WT and knockout *MDK*-deleted MCC13 tumor cells following IFNβ stimulation. This study confirmed enhanced IFN-response signatures, and additionally suggested enhanced NF-kB, inflammatory, and STAT3 signaling as well as decreased hypoxia signaling in MDK-knockout (KO) compared to control tumor cells (**Fig. 4f**).

MDK-knockout tumor cells demonstrated increased expression of immunomodulatory factors including *C3*, *CXCL6*, *IL6*, *CASP1* and decreased expression of *TGFB1*, among others (**Fig. S4c**). Given the potent and broad effects of *MDK* deletion on IFN-response and NF-kB signaling, we hypothesized that MDK may additionally limit MHC-I expression in neuroendocrine tumors with high MDK expression. We therefore tested MHC-I expression of our MDK-KO MCC cell lines. In agreement with our hypothesis, the loss of MDK enhanced the expression of MHC-I following IFNβ stimulation in both MCC13 and WAGA (**Fig 4g, S4d**. To test whether this key effect also applied to SCLC, we deleted MDK from the SCLC cell lines, DMS153 and DMS79.

We again tested the expression of MHC-I with and without IFNβ stimulation. In this case, we observed significantly greater MHC-I expression both at baseline and following IFN-stimulation in MDK-KO cells compared to controls (**Fig 4h**). These results confirmed that MDK acts to suppress both IFN response and MHC-I expression in a tumor cell-intrinsic manner in patient-derived neuroendocrine tumor cells.

### Effects of MDK signaling on human antigen-presenting cells

Our PERCEPT results from non-responsive MCC tumor samples suggested that tumor and immune cells concordantly failed to respond to IFN and SLR14 stimulation in MDK-high samples. We consequently hypothesized that tumor-derived midkine can suppress antigen-presenting cell activation by these stimulations. Accordingly, we observed CD14+ monocytes co-cultured with MDK-WT tumor cells had decreased upregulation of antigen expression by HLA-A,B,C. Given that IFN signaling is central to antigen-presenting cell (APC) activation, we hypothesized that MDK may additionally suppress this process^13^. To test this hypothesis, we incubated MDK-KO or WT tumor cells with either CD14+ monocytes isolated from healthy donor PBMCs, or with monocyte-derived dendritic cells (moDCs), developed by exposure of monocytes to GM-CSF and IL-4. Whereas co-incubation with MDK-WT MCC cell lines suppressed both MHC-I expression and CD86 in monocytes, this suppression was relieved by deletion of MDK (**Fig. 5a, b, S5a**). We observed a similar effect on monocyte-derived dendritic cells (moDCs) from PBMCs in attenuation of HLA-A,B,C, CD86, and CD80 (**Fig. 5c, S5c**).

**Fig 5.**
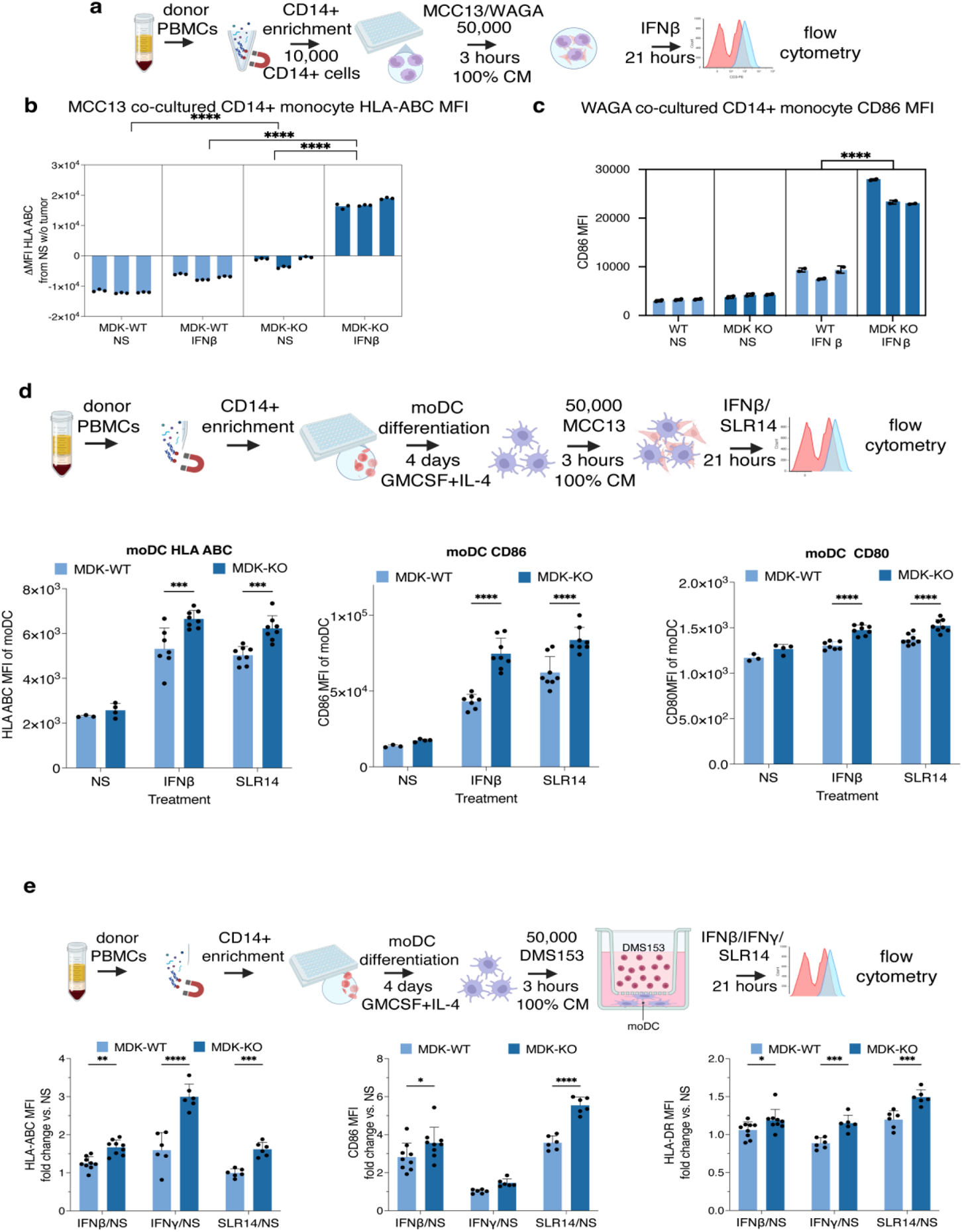
Midkine (MDK) suppresses antigen-presenting cell activation by IFN and SLR14 stimulation. (**a,b**) MCC13 (MCC) or (**a,c**) WAGA (MCC) MDK-WT and MDK-KO tumor cells in 100% conditioned media (CM) co-cultured with CD14+ monocytes magnetically enriched from PBMCs for three hours prior to treatment with 1000 IU/ml IFNβ vs. NS for 21 hours prior to flow cytometry of HLA A,B,C and CD86 on monocytes (>90% CD45+CD11b+CD3-). (**d**) MCC13 (MCC) tumor cells in 100% conditioned media (CM) co-cultured with moDCs differentiated from CD14+ monocytes magnetically enriched from PBMCs for three hours prior to treatment with 1000 IU/ml IFNβ, 1 ug/ml SLR14 vs. NS for 21 hours prior to flow cytometry of HLA-A,B,C, CD86 and CD80 on moDCs (>90% CD45+CD3-). (**e**) DMS153 (SCLC) tumor cells in 100% conditioned media (CM) co-cultured across a cell impermeable transwell (pore 3 um) with moDCs differentiated from CD14+ monocytes magnetically enriched from PBMCs for three hours prior to treatment with 1000 IU/ml IFNβ, 1000 U/ml IFNγ, 1 ug/ml SLR14 vs. NS for 21 hours prior to flow cytometry of HLA-A,B,C, CD86 and HLA-DR on moDCs (>90% CD45+CD3-). *=p < 0.05, **=p < 0.01, ***=p < 0.005, ****=p < 0.001 by two-way ANOVA with Sidak correction for multiple comparisons. Data representative of three independent experiments.

Furthermore, MDK-WT SCLC tumor cells demonstrated suppression of HLA-A,B,C, CD86 and HLA-DR on moDCs across a cell-impermeable transwell after stimulation with IFNβ, IFNγ, or SLR14 in comparison to MDK-KO (**Fig. 5d, S5d,e**), demonstrating that this effect applies to SCLC as well as MCC, and that it is independent of cell-cell contact.

Given the direct observation of MDK as a suppressor of IFN response, antigen presentation and APC activation in PERCEPT experiments and cell culture models, we hypothesized that midkine expression would be associated with poorer immune infiltration and response to immune checkpoint inhibitors in neuroendocrine cancers. To test this hypothesis, we leveraged a database of 872 SCLC patients with clinically annotated responses to immune checkpoint inhibitors and matched RNA sequencing data. In this dataset, we observed a step-wise decrease in event-free survival (EFS, comprising progression of disease, initiation of a new therapy, or death) based on MDK expression quartile, from a median EFS in the highest MDK expression quartile of 11.4 months to 16.9 months in the lowest MDK expression quartile (log-rank p=0.0082) in SCLC patients treated with immune checkpoint inhibitors (**Fig. 6a**). MDK expression additionally anti-correlated with immune infiltration in this cohort with QuanTIseq performed on the highest 50% (MDK-high) vs. lowest 50% (MDK-low) SCLC patient tumors suggesting lower CD8+ T cell and M1-macrophage, and higher M2-macrophage infiltration (**Fig. 6b, S6a**). Both SCLC (**Fig. 6c, S6b;** independent dataset of 79 samples) and MCC (**Fig. 6d, S6c;** independent dataset of 54 samples) patient tumor RNAseq datasets ranked by MDK expression demonstrated significant anticorrelations with gene signatures of antigen presenting cell activity, MHC-I (i.e. HLA-A,B,C), MHC-II (i.e. HLA-DR) and T cells.

**Fig 6.**
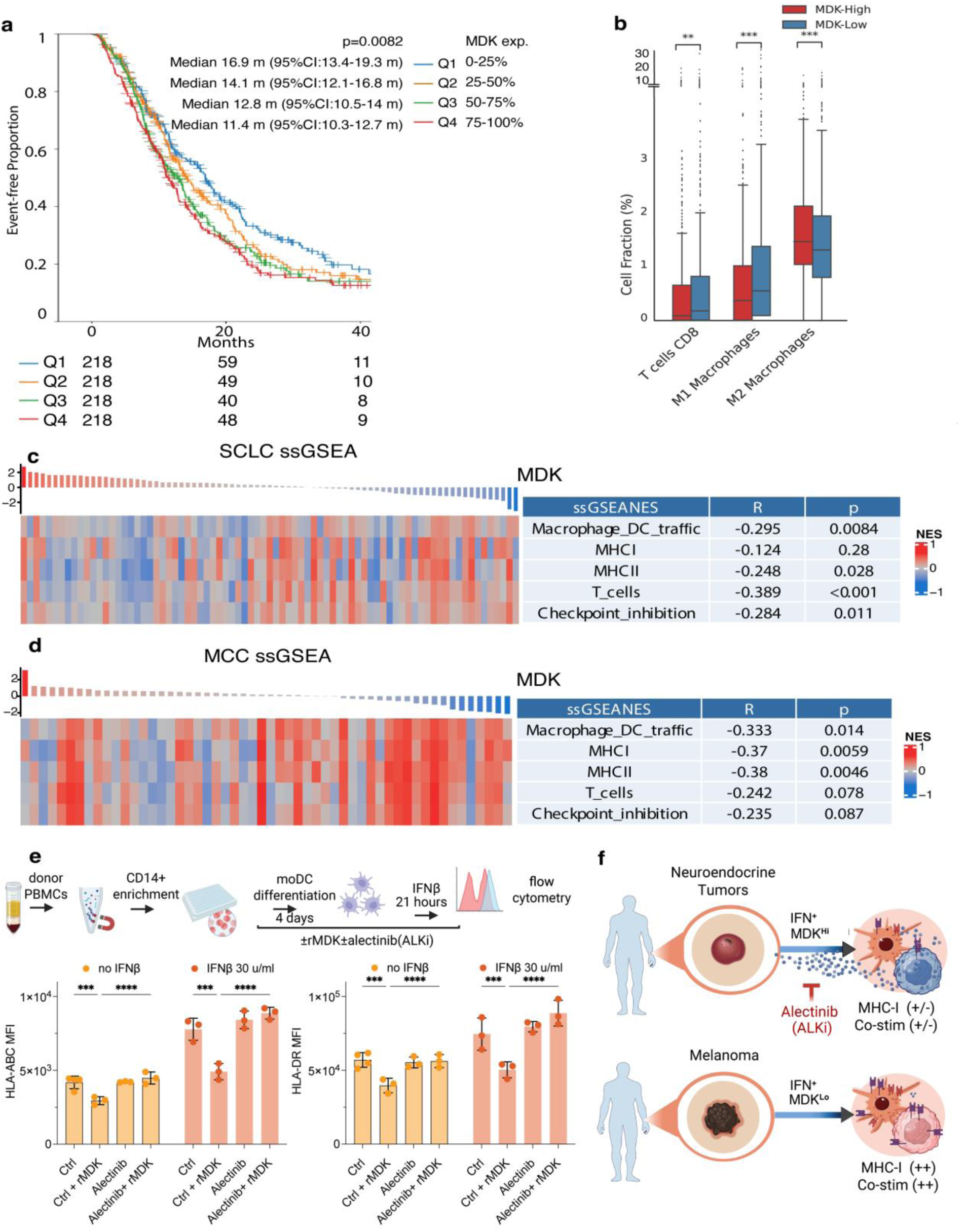
Midkine (MDK) expression is associated with worse event-free survival in SCLC and decreased immune infiltration. (**a**) SCLC patient tumors that underwent comprehensive tumor profiling at Caris Life Sciences were compared by MDK expression quartiles per log-rank test p=0.0082. (**b**) QuanTIseq was performed on the highest 50% (MDK-high) vs. lowest 50% (MDK-low). **=p < 0.01, ***=p < 0.005 by two-way ANOVA with Sidak correction for multiple comparisons. (**c**) ssGSEA of SCLC patient tumor RNAseq ranked by relative MDK expression^31^. (**d**) ssGSEA of MCC patient tumor RNAseq ranked by relative MDK expression^30^. (**e**) Monocyte-derived dendritic cells (moDCs) were differentiated from CD14+ monocytes magnetically enriched from PBMCs with or without rMDK (40 ng/ml) and/or alectinib (1 nM) for 4 days prior to activation with 30 U/ml IFNβ vs. NS for 21 hours prior to flow cytometry of MHC-I (HLA-ABC), MHC-II (HLA-DR) on moDCs (>90% CD45+CD3-). *=p < 0.05, **=p < 0.01, ***=p < 0.005, ****=p < 0.001 by two-way ANOVA with Sidak correction for multiple comparisons. Data representative of three independent experiments. (**f**) Graphical summary of experimental findings that neuroendocrine tumor-derived MDK suppresses MCC and SCLC tumor and DC activation, potentially impeding the tumor immunity cycle at multiple points.

To enable an actionable clinical development path for MDK-inhibiting therapies, we expanded on prior reports of ALKi effects on MDK signaling by testing the effect of the FDA-approved ALK inhibitor, alectinib in tumor cell-DC co-culture assays^28^. We observed that alectinib fully alleviated the MDK-induced inhibition of MHC-I (HLA-A,B,C), MHC-II(HLA-DR), and co-stimulation (CD86) in moDCs even at a 1 nM dose (**Fig. 6e, S7a**). As previously reported, the ALKi NVP-TAE-684 also alleviated MDK-induced inhibition of MHC-I (HLA-A,B,C), MHC-II(HLA-DR) (**Fig. S7b**). ^28^. These results a potential novel application of alecitinib and other ALK inhibitors in MDK-expressing neuroendocrine cancers.

In conclusion, (**Fig. 6f**), midkine, enriched in human neuroendocrine tumor MCC and SCLC patient samples and cell lines, inhibited upregulation of antigen-presentation by tumor cells in response to innate immune stimuli, attenuated monocyte and DC activation of antigen presentation and co-stimulation, and was associated with decreased T cell activation and ICI response.

## Discussion

In this study, we develop and apply PERCEPT, a multiplexed perturbational single-cell RNA sequencing strategy that enables direct testing of therapeutic agents in freshly resected human tumors. By systematically cross-comparing up to ten immunomodulatory perturbations simultaneously in proportionally intact tumor-immune cellular ecosystems, PERCEPT captures lineage- and context-specific programs of innate immune activation and resistance that are inaccessible to conventional model systems. Applying this approach, we identified the RIG-I agonist, SLR14 as a potent inducer of interferon signaling and midkine as a lineage-specific mechanism of resistance to immune signaling in neuroendocrine tumors. These findings establish proof of concept for a framework for causal inference in human immuno-oncology.

PERCEPT experiments build upon the corpus of knowledge gained from existing human single-cell RNAseq studies but overcome the critical barrier of inter-patient heterogeneity by normalizing treatment responses to unstimulated controls^17^. This approach enables the application of causal inference frameworks such as CINEMA-OT and the accurate attribution of response and resistance patterns. Moreover, by comparing numerous treatments, PERCEPT offers the potential to both prioritize treatments and map patient-specific contexts in which each treatment is likely to be effective. As regulatory agencies deprioritize animal models and emphasize the role of generative artificial intelligence in drug development, PERCEPT offers the potential to help fill a key gap in the training data necessary for computational models to predict treatment response and toxicity, mapping the latent spaces of treatment response at high dimensionality and throughput^12^.

Across diverse tumor types, PERCEPT revealed distinct patterns of response to IFN-inducing therapies. We identified a novel RIG-I agonist, SLR14, as capable of inducing robust IFN signaling in tumor cells as well as unique, activated states in T cells. Given the recent promising Phase II clinical trial results with inter-tumoral injection of poly I:C in αPD-1 therapy-resistant melanoma, these data nominate SLR14 as a next-generation innate immune agonist with high translational potential for further investigation^11^. PERCEPT also generated credible predictions of clinical response to immune checkpoint inhibition. Consistent with prior studies, αPD-1 elicited *ex vivo* responses only in tumors not previously refractory to checkpoint blockade.

Intriguingly, however, we observed a response to combined stimulation with αPD-1 and IFNβ in one tumor with clinical resistance to αPD-1^9^. These findings demonstrate that multiplexed perturbational profiling can capture the diversity of patient-specific therapeutic responses and generate experimentally testable hypotheses for the design of rational immuno-oncology combination strategies.

Among our most striking findings, several neuroendocrine tumor samples were almost entirely unresponsive to IFN-inducing therapies and to IFN cytokine exposure. In these samples both tumor and immune compartments failed to respond to treatment, suggesting the potential presence of a cell-extrinsic inhibitory factor. CINEMA-OT analysis of these resistant samples revealed enrichment of midkine, a secreted cytokine previously linked to oncogenic signaling in neuroendocrine tumors and immune evasion in melanoma. Functional validation using CRISPR knockout models in MCC and SCLC confirmed high MDK expression in neuroendocrine cancers and demonstrated that MDK suppresses IFN signaling, antigen presentation and dendritic cell activation. These data suggest potential clinical benefit from midkine inhibition in multiple steps in the tumor immunity cycle, including decreased tumor cell growth, enhanced tumor cell antigen presentation and improved DC function. Our findings in patients are in agreement with recent mechanistic insights from melanoma-focused studies demonstrating midkine suppression of DCs through signaling by ALK/STAT3, but suggest a key difference in the tumor lineages most likely to benefit from midkine inhibition and offer a distinct path for clinical development^27,28^. Indeed, combining our data highlights an opportunity for ALK inhibitors such as alectinib to be investigated in combination with ICB for direct anti-tumor benefits and to overcome a key mechanism of αPD-1 resistance in midkine-high neuroendocrine tumors like MCC and SCLC^28^.

In conclusion, PERCEPT provides a framework for experimental immuno-oncology in patient tumor tissues that enables rigorous, contextual cross-comparison of multiple treatment responses at the level of single-cells, cell types, patients and tumor lineages. By uncovering midkine as a lineage-specific mediator of innate immune response in neuroendocrine tumors and nominating SLR14 as a potent stimulator of IFN response, our study demonstrates how deep, multiplexed profiling of treatment response can uncover actionable mechanisms of tumor immunity. More broadly, PERCEPT enables a path for human-based preclinical modeling that is scalable, translatable, and consistent with emerging priorities for translational drug development and computational biology.

## Methods and materials

### Ethical approval

The study was approved by the Institutional Review Boards at Yale University following Yale Spore in Skin Cancer (IRB protocol # 0609001869). Patients with a surgical resection clinical indication consented to the donation of surgical resected tissue for research use.

### Tumor processing and *ex vivo* culture

Resected human tumors were minced and digested using Native Bacillus licheniformis protease (Creative Enzymes, NATE-0633) on ice for 30 minutes to dissociate into single cell suspension. Tumor single cell suspensions with low viability were further processed using dead cell removal kit (Miltenyi, 130-090-101). Cells were plated and stimulated with 1000U/ml human IFNβ (pbl assay science 11415), 1000 U/ml human IFNγ, (pbl assay science 11500), 500ng/ml polyI:C HMW (invivogen, tlrl-pic) transfected with lipofectamine 2000, 10ug/ml non-transfected poly(I:C) HMW, 2ug/ml SLR14 (gift from the Ana Pyle lab), 1ug/ml ADU-S100 (InvivoGen, tlrl-nacda2r), 10ug/ml anti-human CD279 antibody (Biolegend, 329902), and cultured for up to 48 hours.

PBMCs were isolated by Ficoll (Sigma-Aldrich, F5415) density centrifugation.

### Cell enrichment and 10x sample preparation

Cultured cells were collected stained with TotalSeq anti-human hashtags C0251- C0258 (Biolegend), viability dye (zombie red, Biolegend 423109) and anti-human CD45-FITC (clone HI30, Biolegend 304038) and enriched for live cells and up to 50% immune cells using BD FACS Aria II. Sorted cells were then resuspended to 700-1200 cells per ul and barcoded for multiplexed single cell sequencing using 10x Genomics 5’v2 chemistry (10x Genomics, PN-1000263).

### Sequencing and 10x sample alignment

Single cell RNA sequencing libraries were sequenced on Illumina NovaSeq at read length of 150bp pair end and depth of 300 million reads per sample.

Cellranger software v6.0.1 (10x Genomics) and prior was used to align single cell RNA sequencing reads to the reference genome GRCh38, count gene expression and capture antibody hashtags.

### PERCEPT scRNA-seq and CITE-Seq Preprocessing

Seurat R package was used to perform single cell antibody hashtag demultiplexing, quality control, clustering and visualizations. Specifically, the HTODemux function from Seurat package was used to assign each cell with its corresponding hashtag labeling(s) and based on the assignment, the cell is classified to be either a singlet, doublet or negative. Singlets are kept for further analyses, and they were assigned to their condition of origin. Quality control was done based on nCount_RNA (the total number of molecules detected within a cell), nFeature_RNA (the number of unique genes detected within a cell), percent_mt (percentage of genes traced back to the mitochondrial genome). Normalization, scaling, PCA and unsupervised clustering were also performed using Seurat and cell type annotation for each cluster was performed based on the differentially expressed gene results by FindAllMarkers. Harmony was used to perform batch correction for different samples to create a merged object. UMAP was performed and used as the primary visualization projection.

### MELD

MELD was implemented on the preprocessed scRNA-seq data as previously described. The relative likelihood ratios (MELD score) were calculated for each pair-wise comparisons between the non-stim and each experimental perturbation. Then within each comparison, the mean MELD score for each sample is calculated. Based on that, the samples were defined as Responder (R) or non-responder (NR) for that perturbation.

### CINEMA-OT

CINEMA-OT was implemented on the preprocessed scRNAseq as previously described^17^. Based on MELD score to SLR14 and IFNβ, melanoma and MCC samples were defined as Responder (R) or non-responder (NR). In CINEMA-OT treatment effect space, genes significantly associated with NR were ranked by T-score.

### CellChat

Annotated cells from melanoma and MCC patients were analyzed separately for CellChat using human Midkine pathway in CellChatDB with the addition of MDK-NOTCH2 interaction^32^. The top 4% of interactions were kept for clarity and simplicity.

### RNA-seq

Bulk RNA sequence reads from NIH dbGap (study accession phs002260.v2.p1) under authorized access were trimmed using fastp (v0.23.2) to remove adaptor sequences. Salmon (v1.4.0) was used to map and quantify the reads to the reference human genome (GRCh38.p14 primary assembly). Transcripts were imported into R and summarized to the gene level with tximport. Ensembl gene IDs were retained as primary identifiers and mapped to HGNC symbols. Differential expression was performed with DESeq2, and low-count genes (<10) were removed across all samples. Volcano plots and heatmaps were generated with ggplot2/ComplexHeatmap. The comparison for gene expression across different datasets was performed after combining and normalizing them together.

### Life Science Cohort

To provide a broad description of expression levels of MDK across tumor types, we gathered data from a large historical cohort sequenced at Caris Life Sciences. Clinical data were acquired from insurance claims, and the selection of systemic therapies was at the discretion of the treating physician.

Whole transcriptome sequencing (WTS) was performed on formalin-fixed paraffin-embedded (FFPE) tumor samples using a hybrid-capture method with the Agilent SureSelect Human All Exon V7 bait panel (Agilent Technologies; RRID) and sequenced on the Illumina NovaSeq platform. Pathologic review was conducted to confirm at least 20% tumor content and adequate tumor size for microdissection and enrichment of tumor-specific RNA. RNA was extracted with the Qiagen FFPE Tissue Kit and assessed for quality and quantity using the Agilent TapeStation. Biotinylated RNA probes were hybridized to cDNA targets, followed by post-capture PCR to generate sequencing libraries, which were quantified, pooled, and sequenced. Demultiplexing was performed using the Illumina DRAGEN FFPE accelerator. Reads were aligned to GRCh37/hg19 using the STAR aligner (v2.7.4a), and transcript quantification was completed with Salmon, yielding expression data for 22,948 genes. BAM files from STAR were processed through a proprietary pipeline for RNA variant detection. The assay demonstrated ≥97% positive percent agreement, ≥99% negative percent agreement, and ≥99% overall agreement with a validated comparator.

We estimated the proportions of ten immune cell types in the tumor microenvironment using QuanTIseq, a computational tool (RRID:SCR_022993) that analyzes RNA expression profiles. QuanTIseq’s accuracy for quantifying myeloid dendritic cells, regulatory T cells, CD8+ and CD4+ T cells, natural killer cells, neutrophils, monocytes, M1 and M2 macrophages, and B cells has been validated against flow cytometry and immunohistochemistry.

### Gene-Set Enrichment Analysis

Variance-stabilized log₂ counts (baseMean > 50) from DESseq2 were ranked by adjusted p-value and served as the pre-ranked statistics. MSigDB Hallmark and Canonical (C2) gene sets (v7.5.1) were used. Enrichment was computed with clusterProfiler (4.12.0) and fgsea (v1.30.0). Normalized enrichment score (NES) and Benjamini–Hochberg FDR < 0.05 were used for significance.

### Cytosig

Normalized gene-expression values were used as input for cytokine-activity scoring with Cytosig. For each sample, Cytosig calculates a normalized enrichment score (NES) for 43 cytokine/chemokine signaling pathways by comparing the ranked sample expression profile to curated up- and down-regulated gene sets.

### VIPER

Activities were inferred with decoupleR (v 2.14.0) on the log-transformed VST matrix. Differential TF activity between SLR14 and IFNβ cells was assessed with FindMarkers(). Volcano plots were rendered with EnhancedVolcano().

### SLR14 priming of T cell responses

100,000 PBMCs per ml RPMI from healthy donors were transfected with lipofectamine 2000 (ThermoFisher) with or without 2ug/ml SLR14 for 48 hours prior to stimulation with aCD3/28 (StemCell) and Brefeldin A (BD biosciences). Intracellular cytokine staining was performed as previously described.

### MDK Crispr KO generation

Nucleofector (Lonza).

Targets all isoforms of MDK at exon 1.

Ko1MDK: TTTCTTTTTGGCGACCGCGG. Cut position 46382116. Ko2MDK: CGGGGAGCGAGTGCGCTGAG. Cut position 46382113

For CRISPR-mediated knockout, MCC13, WAGA, DMS79 and DMS153 were subjected to electroporation using Lonza 4D-Nucleofector with the SE Cell Line 4D-Nucleofector Kit. Cells were prepared according to the manufacturer’s instructions and resuspended in supplemented electroporation buffer. Ribonucleoprotein (RNP) complex containing Cas9 protein, and guide RNA (gRNA) was used for each electroporation.

The gRNA sequences targeting MDK were designed using benchling and synthesized by Integrated DNA Technologies (IDT). The gRNA sequences were as follows:

1. gRNA1: 5’-TTTCTTTTTGGCGACCGCGG-3’
2. gRNA2: 5’-CGGGGAGCGAGTGCGCTGAG-3’

Electroporation was performed under optimized settings for each cell line, and cells were transferred immediately to pre-warmed complete culture medium for recovery at 37°C in a 5% CO2 incubator. Post-electroporation, cells were plated and expanded for initial recovery. Single-cell clones were generated by limiting dilution into 96-well plates and cultured until clonal colonies formed. Genomic DNA was extracted from individual clones, and the target locus was amplified by PCR. Editing efficiency and mutations were assessed by sequencing and further analyzed using TIDE (Tracking of Indels by Decomposition).

Clones confirmed to carry the desired edits were expanded and further validated by ELISA (Abcam #ab193761). MDK-WT cells maintained MDK expression after electroporation and single-cell cloning. At least 3 single clones of validated MDK-WT or MDK-KO were pooled for further experimentation.

### Tumor-immune co-culture

MCC (MCC13 or WAGA) or SCLC (DMS153 or DMS79) MDK-WT and MDK-KO tumor cells (10^5^ cells/ml) in 100% conditioned media (CM) were co-cultured with either fresh CD14+ monocytes magnetically enriched from PBMCs (>90% CD45+CD3-) or after moDC differentiation with 250 IU/mL IL-4 and 800 IU/mL GM-CSF (Miltenyi Biotec #130-094-812) for 3 days with moDC morphology confirmed by microscopy. Alectinib (Selleckchem #2762) or NVP-TAE-684 (Selleckchem #S1108) were used at indicated concentrations. rMDK (Peprotech #450-16) was used at 40 nM. As indicated, SCLC tumor cells were co-cultured across a cell-impermeable transwell (pore 3 um, Corning #3399). After three hours of co-culture, cells were stimulated with 1000 IU/ml IFNβ (PBL Assay Science, cat. #11415), 1000 U/ml IFNγ (PBL Assay Science, cat. #11500), 1 ug/ml SLR14 (gift of Pyle lab) vs. NS for 21-24 hours prior to harvest with trypsin (Thermofisher #25200-056).

### Flow cytometry

After treatment, cells were harvested and stained for surface expression of major histocompatibility complex class I molecules HLA-A,B,C (clone W6/32, BioLegend). CD86 (clone GL-1, BioLegend) and HLA-DR (clone LN3, Biolegend) on moDCs (CD45+ (clone HI30, Biolegend) CD3- (clone OKT3, Biolegend) CD11b+ (clone ICRF44, Biolegend)). T cells were CD45+ CD3+ and either CD8+ (clone SK1, Biolegend) or CD4+ (clone SK3, Biolegend) and profiled for Gzmb (clone QA16A02, Biolegend), and IFNγ (clone XMG1.2, Biolegend). Live tumor and immune cells were gated using Zombie Aqua fixable viability dye (BioLegend) to exclude dead cells prior to analysis by flow cytometry using the Cytoflex S running CytExpert 2.4 (all Beckman Coulter). All assays were performed in three independent biological replicates. Results were analyzed by Flowjo v10 (Flowjo) and Prism v10 (GraphPad by Dotmatics).

## Acknowledgments

We thank patients and their families for donating surgical specimens through the Specimen Resource Core of the Yale SPORE in Skin Cancer. PBMCs were prepared by Nicholas Schindler. SLR14 kindly provided by Olga Federova and Anna Marie Pyle. The results published here are in part based upon data generated by the TCGA Research Network: https://www.cancer.gov/tcga. This work was supported by AstraZeneca, 1R37CA279834, 5P50CA121974, 3P50DE030707. C.J.P was supported by T32CA233414, Conquer Cancer - Endowed Young Investigator Award, in Memory of John R. Durant, MD, and 5K12CA215110. L.L was supported by T32AI07019. J.W.A. was supported by T32GM136651. COI: A.I. co-founded RIGImmune, Xanadu Bio and PanV, and is a member of the Board of Directors of Roche Holding and Genentech. A.I. is a co-inventor on patents related to Stem-Loop RNA (SLR) compounds.

## Supplemental Figures

**Fig S1.**
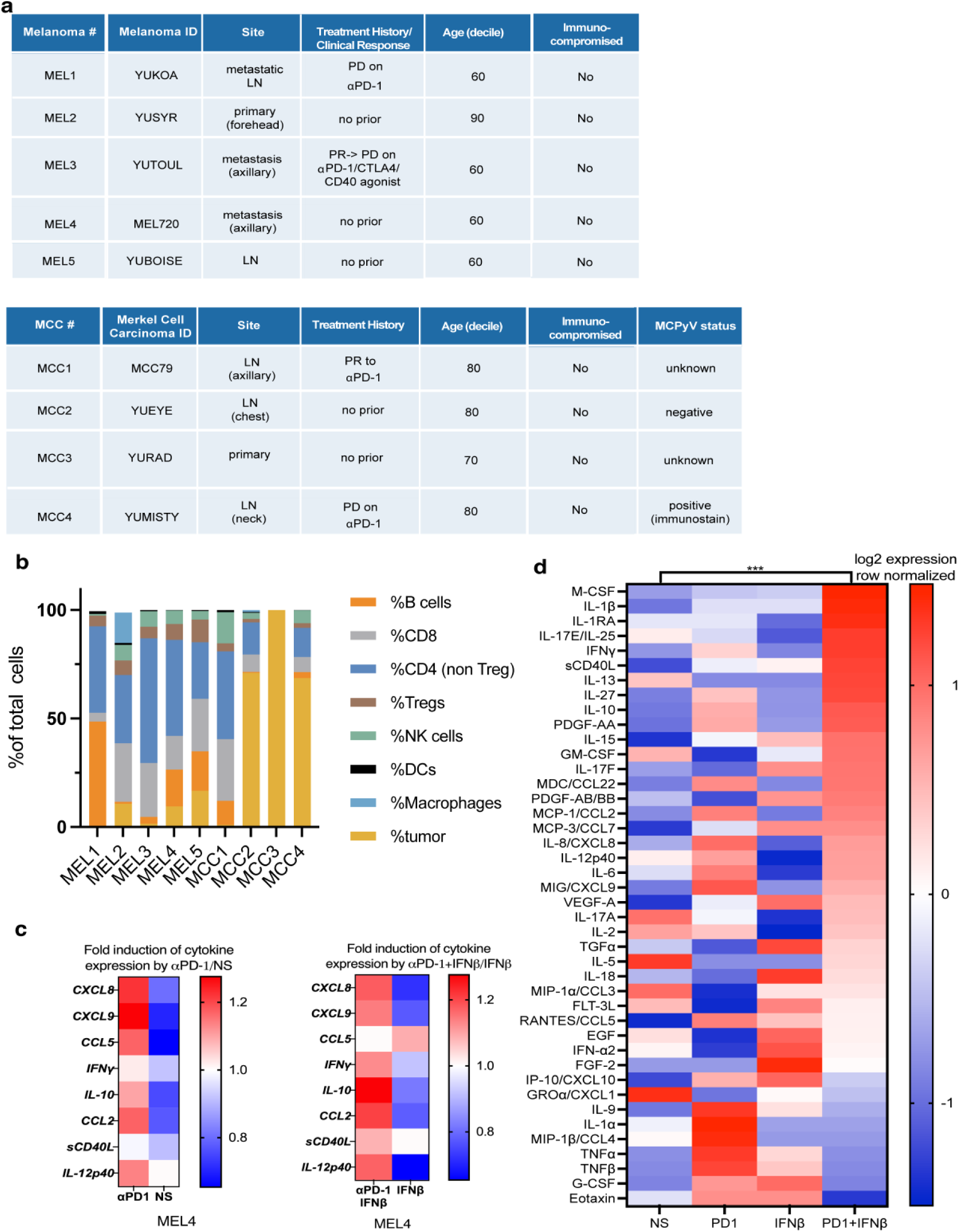
(**a**) Melanoma and MCC patient sample clinical characteristics. (**b**) scRNAseq clusters by cellular identity per sample. (**c**) Eve Technologies serum cytokine concentrations of MEL4 αPD-1/NS and αPD-1+ IFNβ / IFNβ. (**d**) log2 expression row normalized of MEL4 treated with NS, αPD-1, IFNβ, and αPD-1+ IFNβ.

**Fig S2.**
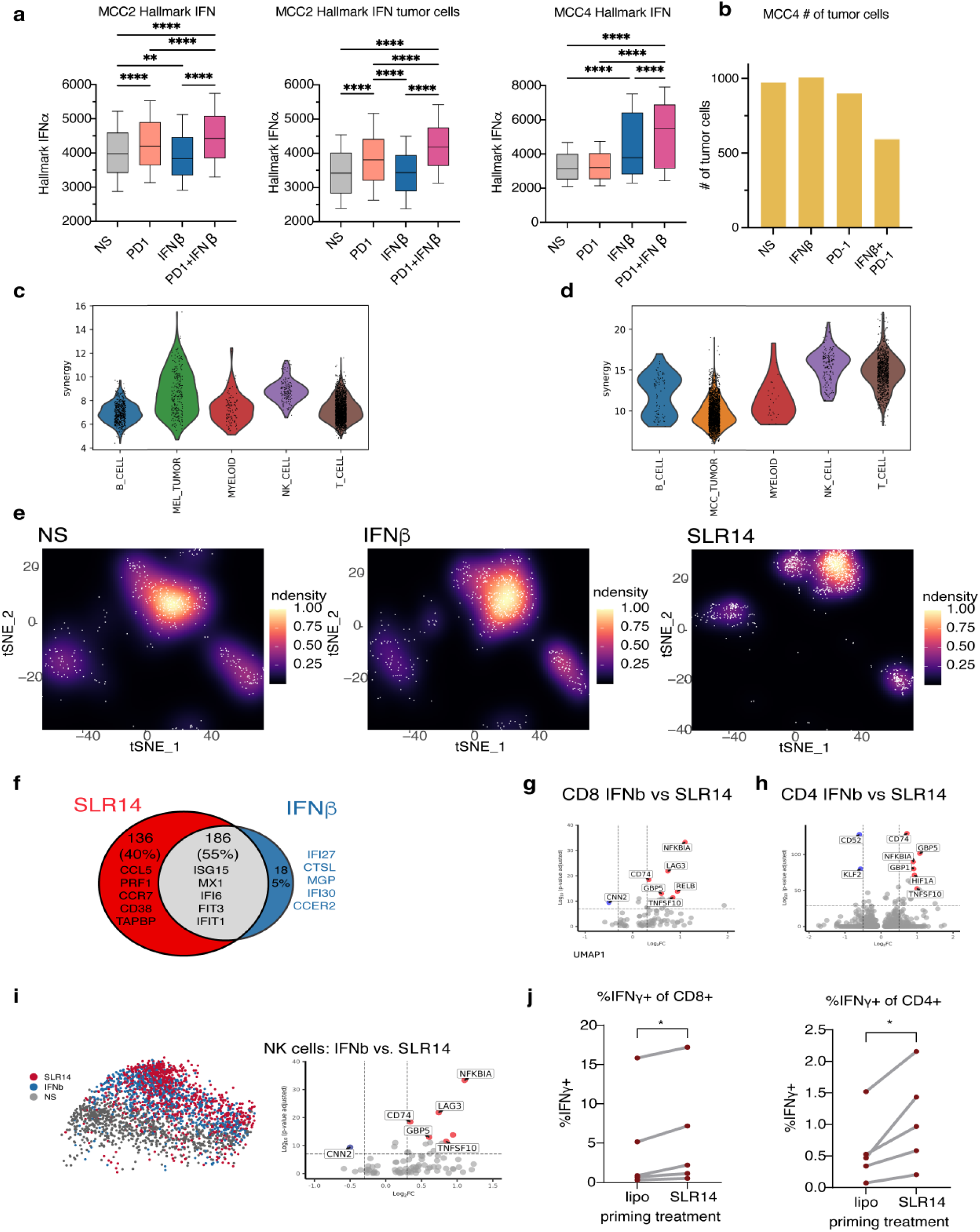
(**a**) Hallmark IFN signature of MCC2 or MCC4 total or tumor cells, respectively. **0.01, ****=p < 0.001 by Mann-Whitney test. (**b**) # of tumor cells by scRNAseq of MCC4. (**c**) Synergy score by CINEMA-OT of scRNAseq from melanoma tumor samples by cell type. (**d**) Synergy score by CINEMA-OT of scRNAseq from MCC tumor samples by cell type. (**e**) t-SNE representation of CD8+ T cells under NS, IFNβ or SLR14. (**f**) scRNAseq differential gene expression of SLR14/NS and IFNβ/NS. (**g**) volcano plot of differential gene expression of CD8 and (**h**) CD4 T cells after SLR14 and IFN. (**i**) UMAP and volcano plot of differential gene expression of NK cells after SLR14 and IFNβ. (**j**) PBMCs were isolated and primed with SLR14 or lipofectamine control for 48 hours as per *ex vivo* tumor protocol, before 6-hour stimulation and αCD3/28 stimulation in Brefeldin A to measure intracellular IFNγ production. P < *0.05, **0.01, from t-test between lipofectamine control and SLR14.

**Fig S3.**
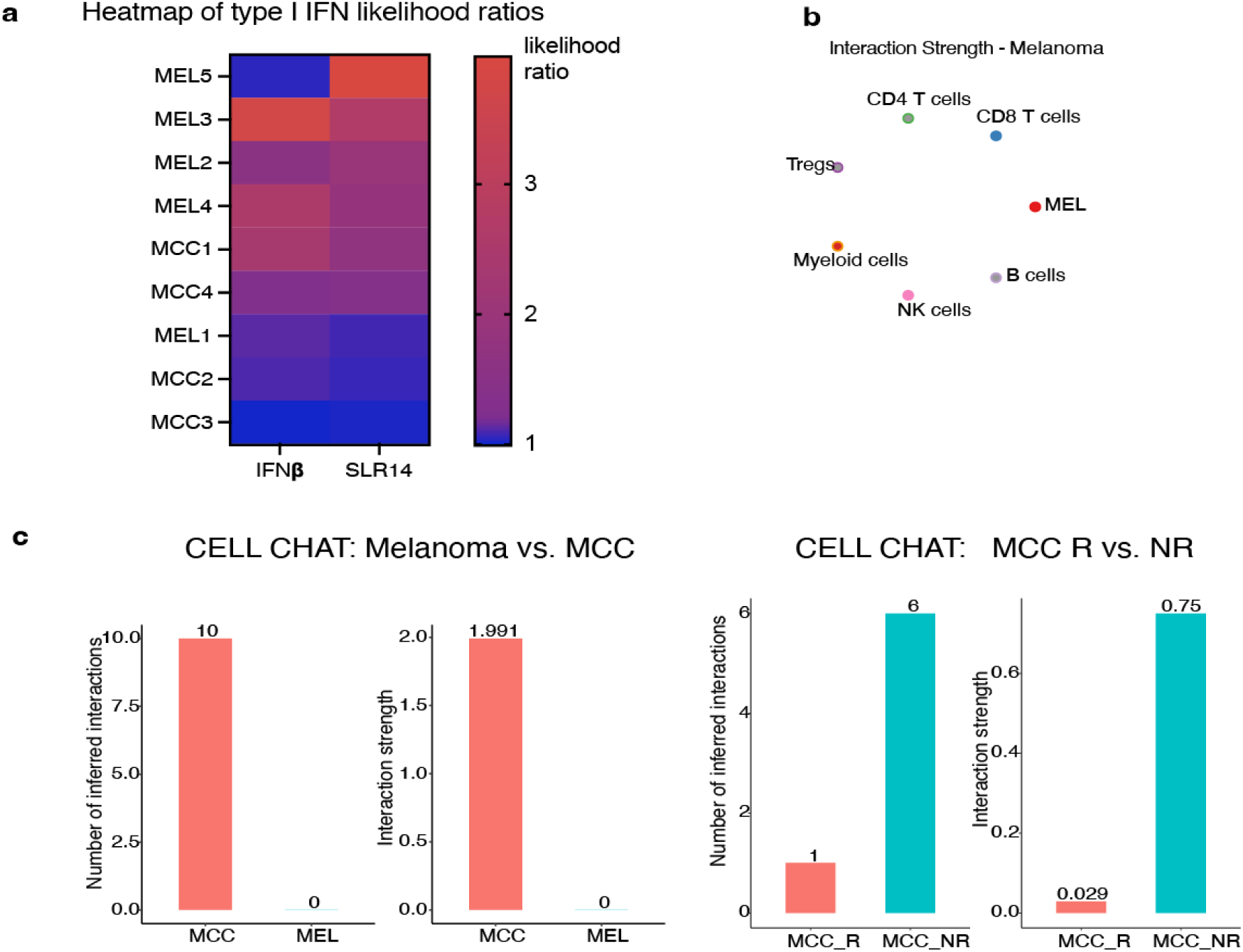
(**a**) PERCEPT samples the likelihood of response to SLR14 or IFNβ by MELD score was measured. (**b**) CellChat analysis of the MDK signaling pathway from melanoma samples by cell type (highest 4%). (**c**) CellChat analysis of the MDK signaling pathway comparing MCC vs. MEL and MCC_Responder vs. MCC_Non-Responder.

**Fig S4.**
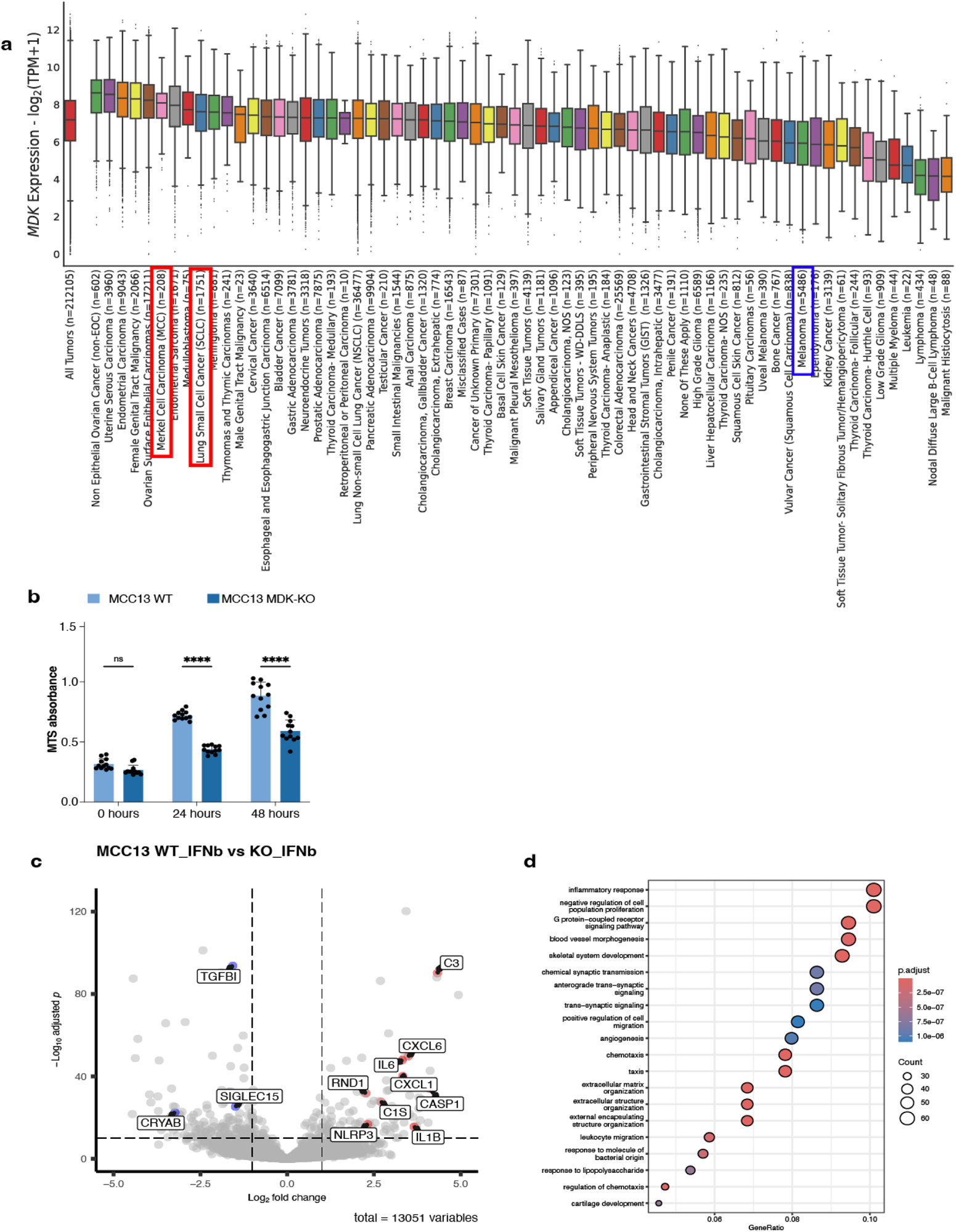
(**a**) Patient tumors that underwent comprehensive tumor profiling at Caris Life Sciences were compared by MDK expression. (**b**) 10^5^ MCC13 MDK-WT and MDK-KO tumor cells were plated per 96-well plate and MTS assay performed at indicated times after plating. (**c**) Volcano plot of differential gene expression and ssGSEA of RNAseq of MCC13 MDK-WT and MDK-KO treated with 1000 IU/ml IFNβ for 24 hours. (**d**) MCC13 (MCC) MDK-WT and MDK-KO tumor cells treated with 1000 IU/ml IFNβ, 1000 U/ml IFNγ, or NS for 24 hours prior to flow cytometry of HLA A,B,C on live cells. ****=p < 0.001 by two-way ANOVA with Sidak correction for multiple comparisons. Data representative of three independent experiments.

**Fig S5.**
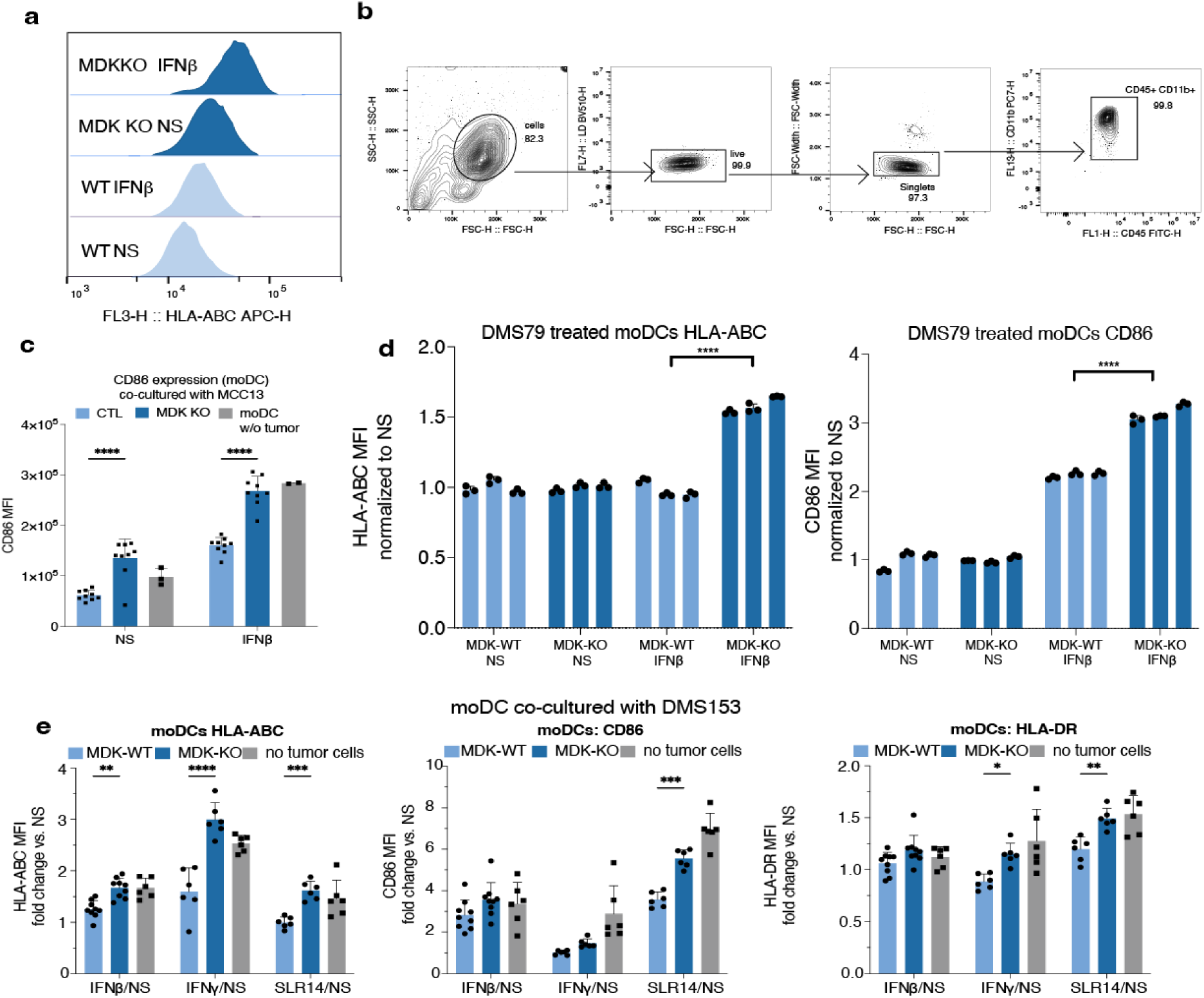
(**a**) MCC13 (MCC) MDK-WT and MDK-KO tumor cells treated with 1000 IU/ml IFNβ, 1000 U/ml IFNγ, or NS for 24 hours prior to flow cytometry of HLA A,B,C on live cells. ****=p < 0.001 by two-way ANOVA with Sidak correction for multiple comparisons. (**b**) MCC13 (MCC) MDK-WT and MDK-KO tumor cells in 100% conditioned media (CM) co-cultured with CD14+ monocytes magnetically enriched from PBMCs for three hours prior to treatment with 1000 IU/ml IFNβ vs. NS for 21 hours prior to flow cytometry of HLA A,B,C (>90% CD45+CD11b+CD3-). (**c**) MCC13 (MCC) tumor cells in 100% conditioned media (CM) co-cultured with moDCs differentiated from CD14+ monocytes magnetically enriched from PBMCs for three hours prior to treatment with 1000 IU/ml IFNβ, 1 ug/ml SLR14 vs. NS for 21 hours prior to flow cytometry of HLA-A,B,C, CD86 and CD80 on moDCs (>90% CD45+CD3-). (**d**) DMS79 (SCLC) tumor cells in 100% conditioned media (CM) co-cultured across a cell impermeable transwell (pore 3 um) with moDCs differentiated from CD14+ monocytes magnetically enriched from PBMCs for three hours prior to treatment with 1000 IU/ml IFNβ vs. NS for 21 hours prior to flow cytometry of HLA-A,B,C and CD86 on moDCs (>90% CD45+CD3-). ****=p < 0.001 by two-way ANOVA with Sidak correction for multiple comparisons. Data representative of three independent experiments. (**e**) DMS153 (SCLC) tumor cells in 100% conditioned media (CM) co-cultured across a cell impermeable transwell (pore 3 um) with moDCs differentiated from CD14+ monocytes magnetically enriched from PBMCs for three hours prior to treatment with 1000 IU/ml IFNβ, 1000 U/ml IFNγ, 1 ug/ml SLR14 vs. NS for 21 hours prior to flow cytometry of HLA-A,B,C, CD86 and HLA-DR on moDCs (>90% CD45+CD3-). *=p < 0.05, **=p < 0.01, ***=p < 0.005, ****=p < 0.001 by two-way ANOVA with Sidak correction for multiple comparisons. Data representative of three independent experiments.

**Fig S6.**
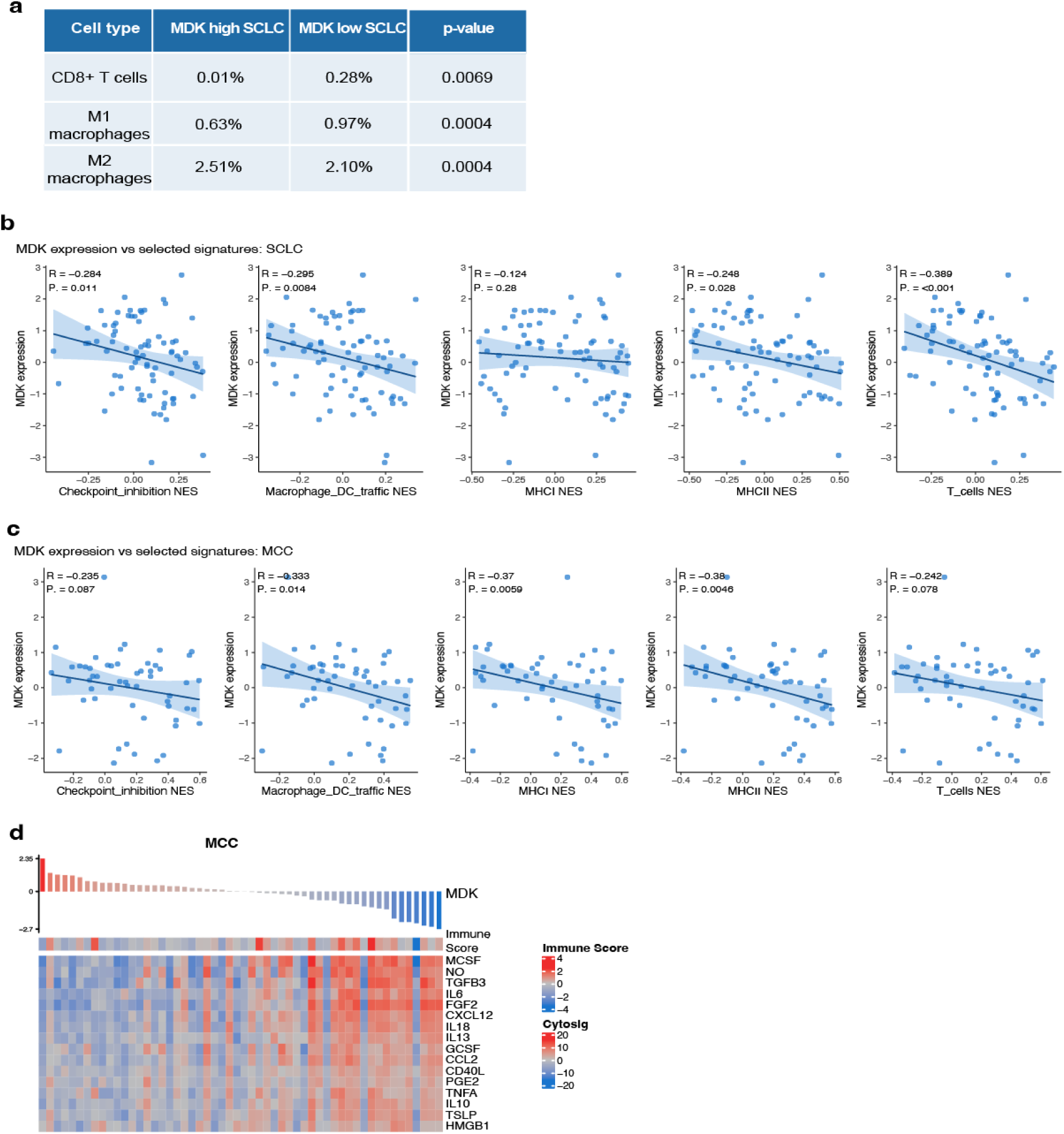
(**a**) QuanTIseq was performed on the SCLC tumors with highest 50% (MDK-high) vs. lowest 50% (MDK-low). **=p < 0.01, ***=p < 0.005 by two-way ANOVA with Sidak correction for multiple comparisons. (**b**) Spearman’s rank correlation coefficients of normalized enrichment score (NES) by MDK expression in patient tumors with SCLC. **(c**) Spearman’s rank correlation coefficients of normalized enrichment score (NES) by MDK expression in patient tumors with MCC. (**d**) Immune gene expression related to ssGSEA of MCC patient tumor RNAseq ranked by relative MDK expression.

**Fig S7.**
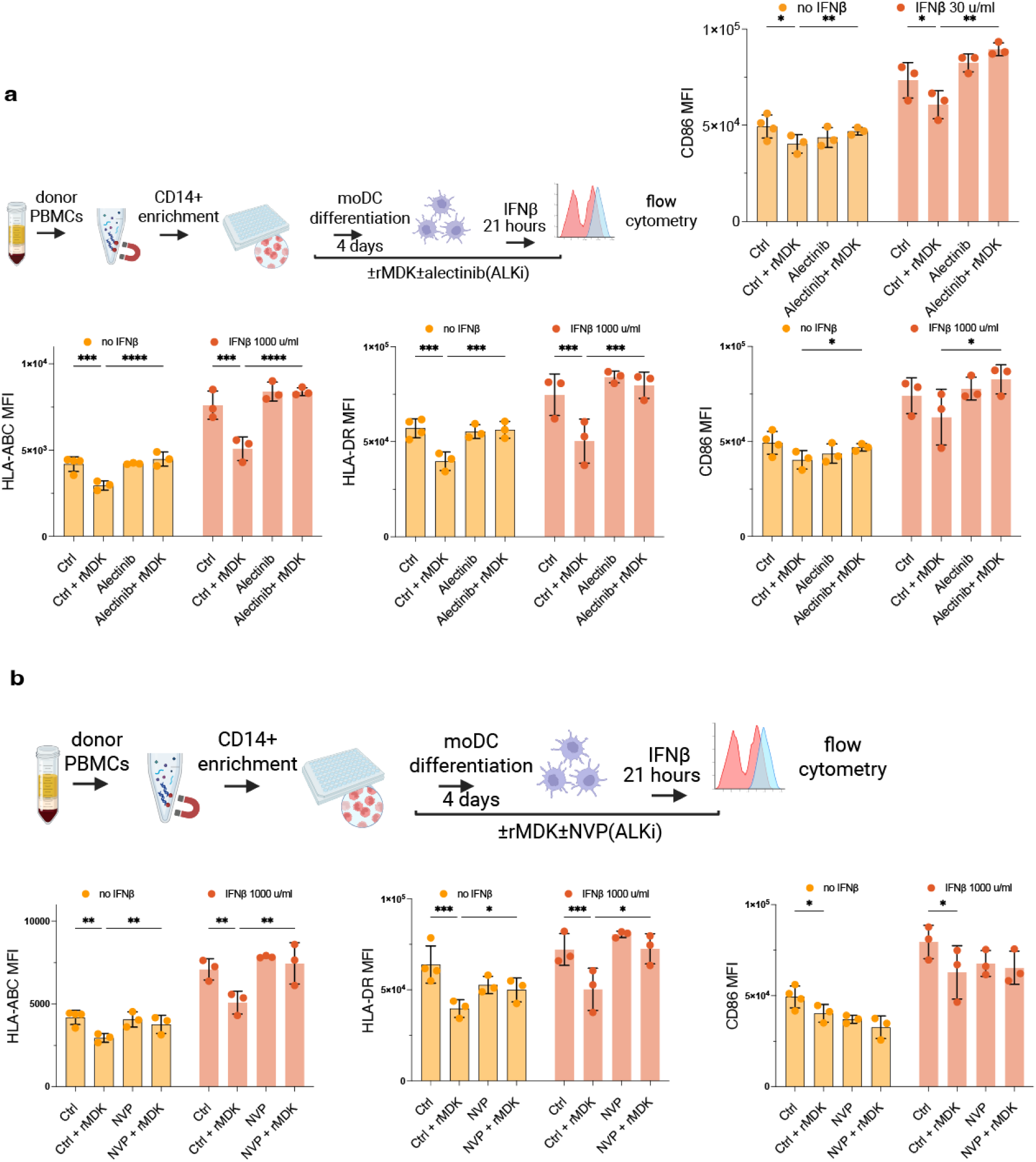
(**a**) Monocyte-derived dendritic cells (moDCs) were differentiated from CD14+ monocytes magnetically enriched from PBMCs with or without rMDK (40 ng/ml) and/or alectinib (1 nM) for 4 days prior to activation with 30 or 1000 U/ml IFNβ vs. NS for 21 hours prior to flow cytometry of MHC-I (HLA-ABC), MHC-II (HLA-DR) or costimulation (CD86) on moDCs (>90% CD45+CD3-). (**b**) Monocyte-derived dendritic cells (moDCs) were differentiated from CD14+ monocytes magnetically enriched from PBMCs with or without rMDK (40 ng/ml) and/or NVP-TAE-684 (100 nM) for 4 days prior to activation with 1000 U/ml IFNβ vs. NS for 21 hours prior to flow cytometry of MHC-I (HLA-ABC), MHC-II (HLA-DR) or costimulation (CD86) on moDCs (>90% CD45+CD3-). *=p < 0.05, **=p < 0.01, ***=p < 0.005, ****=p < 0.001 by two-way ANOVA with Sidak correction for multiple comparisons. Data representative of three independent experiments.

## References

1. Ehrhardt, S., Appel, L. J. & Meinert, C. L. Trends in National Institutes of Health Funding for Clinical Trials Registered in ClinicalTrials.gov. JAMA 314, 2566 (2015).

2. Nakhoda, S. K. & Olszanski, A. J. Addressing Recent Failures in Immuno-Oncology Trials to Guide Novel Immunotherapeutic Treatment Strategies. Pharm Med 34, 83–91 (2020).

3. Bareham, B., Georgakopoulos, N., Matas-Céspedes, A., Curran, M. & Saeb-Parsy, K. Modeling human tumor-immune environments in vivo for the preclinical assessment of immunotherapies. Cancer Immunol Immunother 70, 2737–2750 (2021).

4. Daei Farshchi Adli, A., Jahanban-Esfahlan, R., Seidi, K., Samandari-Rad, S. & Zarghami, N. An overview on Vadimezan (DMXAA): The vascular disrupting agent. Chem Biol Drug Des 91, 996–1006 (2018).

5. Majumder, B. et al. Predicting clinical response to anticancer drugs using an ex vivo platform that captures tumour heterogeneity. Nat Commun 6, 6169 (2015).

6. Jenkins, R. W. et al. *Ex Vivo* Profiling of PD-1 Blockade Using Organotypic Tumor Spheroids. Cancer Discovery 8, 196–215 (2018).

7. Sivakumar, R. et al. Organotypic tumor slice cultures provide a versatile platform for immuno-oncology and drug discovery. OncoImmunology 8, e1670019 (2019).

8. Roelants, C. et al. Ex-Vivo Treatment of Tumor Tissue Slices as a Predictive Preclinical Method to Evaluate Targeted Therapies for Patients with Renal Carcinoma. Cancers 12, 232 (2020).

9. Voabil, P. et al. An ex vivo tumor fragment platform to dissect response to PD-1 blockade in cancer. Nat Med 27, 1250–1261 (2021).

10. Basak, N. P. et al. Tumor histoculture captures the dynamic interactions between tumor and immune components in response to anti-PD1 in head and neck cancer. Nat Commun 15, 1585 (2024).

11. Márquez-Rodas, I. et al. BO-112 Plus Pembrolizumab for Patients With Anti–PD-1–Resistant Advanced Melanoma: Phase II Clinical Trial SPOTLIGHT-203. JCO (2025) doi:10.1200/jco-24-02595.

12. FDA. Roadmap to Reducing Animal Testing in Preclinical Safety Studies. (2025).

13. Zhao, H. et al. Inflammation and tumor progression: signaling pathways and targeted intervention. Sig Transduct Target Ther 6, 263 (2021).

14. Iurescia, S., Fioretti, D. & Rinaldi, M. The Innate Immune Signalling Pathways: Turning RIG-I Sensor Activation against Cancer. Cancers 12, 3158 (2020).

15. Lin, Y. et al. New insights on anti-tumor immunity of CD8+ T cells: cancer stem cells, tumor immune microenvironment and immunotherapy. J Transl Med 23, 341 (2025).

16. Márquez-Rodas, I. et al. BO-112 Plus Pembrolizumab for Patients With Anti–PD-1–Resistant Advanced Melanoma: Phase II Clinical Trial SPOTLIGHT-203. JCO 43, 2806–2815 (2025).

17. Dong, M. et al. Causal identification of single-cell experimental perturbation effects with CINEMA-OT. Nat Methods 20, 1769–1779 (2023).

18. Lim, C. S. et al. TLR3 forms a highly organized cluster when bound to a poly(I:C) RNA ligand. Nat Commun 13, 6876 (2022).

19. Burkhardt, D. B. et al. Quantifying the effect of experimental perturbations at single-cell resolution. Nat Biotechnol 39, 619–629 (2021).

20. Philip, M. & Schietinger, A. CD8+ T cell differentiation and dysfunction in cancer. Nat Rev Immunol 22, 209–223 (2022).

21. McLane, L. M., Abdel-Hakeem, M. S. & Wherry, E. J. CD8 T Cell Exhaustion During Chronic Viral Infection and Cancer. Annu. Rev. Immunol. 37, 457–495 (2019).

22. Combes, A. J. et al. Discovering dominant tumor immune archetypes in a pan-cancer census. Cell 185, 184–203.e19 (2022).

23. Street, K. et al. Slingshot: cell lineage and pseudotime inference for single-cell transcriptomics. BMC Genomics 19, (2018).

24. Miller, B. C. et al. Subsets of exhausted CD8+ T cells differentially mediate tumor control and respond to checkpoint blockade. Nat Immunol 20, 326–336 (2019).

25. Jin, S., Plikus, M. V. & Nie, Q. CellChat for systematic analysis of cell–cell communication from single-cell transcriptomics. Nat Protoc 20, 180–219 (2025).

26. Filippou, P. S., Karagiannis, G. S. & Constantinidou, A. Midkine (MDK) growth factor: a key player in cancer progression and a promising therapeutic target. Oncogene 39, 2040–2054 (2020).

27. Cerezo-Wallis, D. et al. Midkine rewires the melanoma microenvironment toward a tolerogenic and immune-resistant state. Nat Med 26, 1865–1877 (2020).

28. Catena, X. et al. Systemic rewiring of dendritic cells by melanoma-secreted midkine impairs immune surveillance and response to immune checkpoint blockade. Nat Cancer 6, 682–701 (2025).

29. The Cancer Genome Atlas Research Network et al. The Cancer Genome Atlas Pan-Cancer analysis project. Nat Genet 45, 1113–1120 (2013).

30. Knepper, T. C. et al. The Genomic Landscape of Merkel Cell Carcinoma and Clinicogenomic Biomarkers of Response to Immune Checkpoint Inhibitor Therapy. Clinical Cancer Research 25, 5961–5971 (2019).

31. Chan, J. M. et al. Signatures of plasticity, metastasis, and immunosuppression in an atlas of human small cell lung cancer. Cancer Cell 39, 1479–1496.e18 (2021).

32. Kishida, S. et al. Midkine Promotes Neuroblastoma through Notch2 Signaling. Cancer Research 73, 1318–1327 (2013).

